# Structure and efflux mechanism of the yeast pleiotropic drug resistance transporter Pdr5

**DOI:** 10.1101/2021.02.09.430491

**Authors:** Andrzej Harris, Manuel Wagner, Dijun Du, Stefanie Raschka, Holger Gohlke, Sander H. J. Smits, Ben F. Luisi, Lutz Schmitt

**Affiliations:** Department of Biochemistry, University of Cambridge; 80 Tennis Court Road, Cambridge, CB2 1GA, United Kingdom; Institute of Biochemistry, Heinrich Heine University Düsseldorf; Universitätsstraße 1, 40225 Düsseldorf, Germany; Institute of Pharmaceutical and Medicinal Pharmacy, Heinrich Heine University Düsseldorf; Universitätsstraße 1, 40225 Düsseldorf, Germany; Center for Structural Studies, Heinrich Heine University Düsseldorf; Universitätsstraße 1, 40225 Düsseldorf, Germany

## Abstract

Pdr5, a member of the extensive ABC transporter superfamily, is representative of a clinically relevant subgroup involved in pleiotropic drug resistance. Through the coupling of nucleotide hydrolysis with drug efflux, Pdr5 homologues enable pathogenic species to survive in the presence of chemically diverse antifungal agents. Our structural and functional results reveal details of an ATP-driven conformational cycle, which mechanically drives drug translocation through an amphipathic channel, and a clamping switch within a conserved linker loop that acts as a nucleotide sensor. One half of the transporter remains nearly invariant throughout the cycle, while its partner undergoes changes that are transmitted across interdomain interfaces to support a peristaltic motion of the pumped molecule. The efflux model proposed here rationalises the pleiotropic impact of Pdr5 and opens avenues for the development of effective antifungal compounds.

## Results and discussion

### Introduction

The ATP-binding cassette (ABC) transporters comprise an extensive family of membrane transport proteins. Found in all domains of life, these machines are the primary source of active transport across the cell membrane and share a common architecture of two pairs of domains: the transmembrane (TMD) and the nucleotide-binding domain (NBD) (*1*). ATP binding and hydrolysis by the NBDs energises the transport cycle by driving conformational changes in the TMDs that enable substrate passage across the bilayer.

ABC transporters recognise a wide range of substrates, including nutrients and cell-wall components (*2, 3*). Many of the transporters export toxic compounds from cells; indeed, overexpression of these proteins is a key contributor to multi-drug resistance (MDR) phenotypes. In plant and fungi, several efflux pumps involved in MDR are found within the pleiotropic drug resistance (PDR) family, whose members confer resistance to structurally and functionally unrelated drugs and xenobiotics (*2*). The distinguishing architectural features of the PDR family include a structural repeat in which the NBD domain is at the N-terminusof both pseudo-protomers, resulting in a reverse transmembrane topology in comparison with other ABC transporters. PDR proteins also harbour a characteristic N-terminal extension of approximately 160-170 amino acid residues of unknown function (*3*).

Since its discovery 30 years ago (*4*), the ABC transporter Pdr5 from *Saccharomyces cerevisiae* has become an established and widely studied model for PDR proteins in fungi that include major pathogens (*5*). Whilst its physiological substrate (or substrates) is not known, Pdr5 was shown to transport a wide variety of chemicals, including azoles, ionophores, antibiotics and other xenobiotics (*6, 7*). Pdr5 homologs in *Candida albicans* contribute to increased mortality in immune-compromised patients (*8, 9*) and fungicide resistance of the plant pathogen *Botrytis cinerea* (*5, 10*).

ABC transporters share several signature motifs in the NBDs (*11*). Both NBDs contribute residues of these conserved motifs to form two nucleotide-binding sites (NBSs) that bind and hydrolyse ATP. In Pdr5, one of the NBS is catalytically active (NBS2), while the other (NBS1) is inactive due to multiple substitutions in crucial residues of all but one motif. While this asymmetry is shared with many other ABC transporters (e.g., CFTR) (*12*), the PDR subfamily represents the most extreme case (*13*). It is unclear how ATPase-deficiency at this nucleotide-binding site supports the transport process.

Despite the three decades of extensive study of Pdr5 biochemistry, the protein has evaded structural characterisation, leaving a wealth of transport data mostly untapped. Combining improvements in homogenous preparations of Pdr5 (*14*) and advances in single-particle electron cryo-microscopy (cryo-EM) of membrane proteins (*15*), we have been able to elucidate the molecular structure of Pdr5 and obtained insight into its distinctive mechanism.

### Pdr5 reconstituted in a detergent-free system can be imaged with cryo-EM at high-resolution

The membrane environment is critical for the activity of highly allosteric transporters like Pdr5. Several purification protocols have been developed that sustain membrane proteins in a native membrane-like environment (*16–19*). In this study, we reconstituted Pdr5 in peptidiscs, which are short amphipathic bi-helical peptides (*17*) compatible with single-particle cryo-EM (*20, 21*), after purification from *S. cerevisiae* cell membranes. This yielded homogenous particles readily visualised by electron cryo-microscopy (Fig. S1*A-B*). We collected four datasets: apo Pdr5, Pdr5 with added ATP, Pdr5 with added ATP and sodium orthovanadate (V_i_), and Pdr5 with ATP and transport substrate rhodamine 6G (R6G) (*22, 23*). V_i_ is an analogue of inorganic phosphate, and a potent inhibitor of ATPases that traps ABC transporters in a transition-state conformation, by mimicking the action of the γ-phosphate during ATP hydrolysis (*24–27*).

The 3D reconstruction of Pdr5 was carried out predominantly in RELION (*28*) and yielded four near-atomic resolution cryo-EM maps. The sample with Pdr5 alone produced a map containing no nucleotide ligands, which we refer to as the apo-Pdr5 state (Fig. S1*C*). The Pdr5 sample with added ATP yielded a map containing the hydrolysed nucleotide ADP in the canonical and ATP in the inactive site, called here ADP-Pdr5 (Fig. S1*E*). The sample with added ATP and R6G produced a map with the nucleotide composition as above, but also with a bound substrate in the transport cavity, called here R6G-Pdr5 (Fig. S1*I*). The orthovanadate dataset contained Pdr5 in a distinct state, showing the vanadate-trapped hydrolytic intermediate and termed AOV-Pdr5, with ATP in the inactive and ADP-VO_4_^3-^ (or V_i_) in the canonical site (Fig. S1*G*). These maps represent the two most distinct conformational states of Pdr5: inward-facing (substrate-receiving) and outward-facing (substrate-releasing).

The resolution of the maps ranged between 2.8 Å and 3.5 Å, calculated using Fourier shell correlation (*29*) (Fig. S1*D, F, H, J*), revealing well resolved side-chains of bulkier amino acid residues (Fig. 1*E-F*). The quality of the maps enabled the majority (90% amino acid residues) of the structure of this 170-kDa transporter to be traced with certainty for all four states. Some of the external loops of the NBD domains had unresolved features, most likely due to flexibility (Fig. S2 and Fig. S3).

**Fig. 1.**
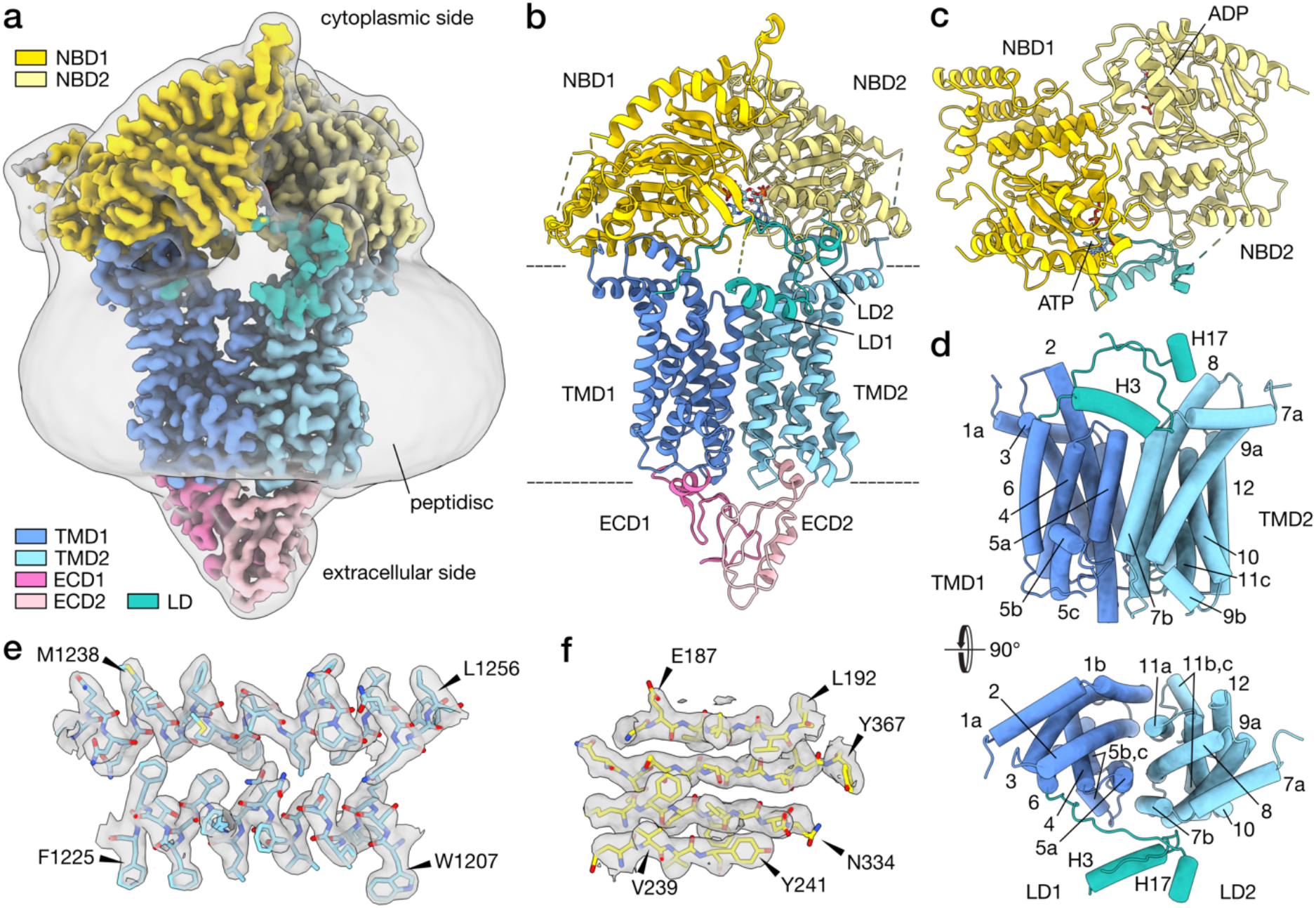
Structural features of Pdr5. (**A**) Depicted here is the cryo-EM map of Pdr5 in inward-open conformation (ADP-Pdr5). The map is coloured according to the domain organisation (see key). The grey transparent envelope surrounding the protein is a Gaussian-filtered (2σ) version of the same map, showing the extent and shape of the peptidisc shell around Pdr5. (**B**) Atomic model of Pdr5 built and refined against the map, in cartoon representation. (**C**) Top view (cytoplasmic side) of the nucleotide-binding domains of Pdr5, showing the position of the bound nucleotides. The linker domain is also depicted. (**D**) Side and top (*Lower*) views of the transmembrane domain of Pdr5 with α-helices shown schematically as cylinders and numbered sequentially (TH1a–12) from the N-terminus (*cf*. Fig. S2 and S3). This panel also includes the linker domain and its two helices H3 and H17. (**E**–**F**) Two portions of the ATP-Pdr5 cryoEM map, showing the reconstructed volume around representative secondary structural features: α-helices TH7b and TH8 of TMD1 domain (E) and a β-sheet in NBD1 domain (F). Arrowheads point to selected amino acid residues. Abbreviations: ECD, extracellular domain; LD, linker domain; NBD, nucleotide-binding domain; TMD, transmembrane domain.

### Pdr5 exhibits the domain architecture of an asymmetric, full-size ABC transporter

The Pdr5 model (Fig. 1*B*) reveals a pseudo-dimeric architecture, undoubtedly the result of gene duplication (*5*). The NBD domains of Pdr5 are structurally similar to those of other members of the superfamily and comprise the RecA-like (*30, 31*) and helical subdomain (the bottom and top halves of NBD1 in Fig. 1*C*). The NBD1 and NBD2 of Pdr5 are structurally similar (Fig. 1*C*), sharing 27% sequence identity. NBD2 is the more conserved of the two; together with the C-loop of NBD1, it forms the composite canonical hydrolytic NBS2, whereas NBD1 and the C-loop of NBD2 cannot sustain ATP hydrolysis as its NBS1 is degenerated (*13, 32*).

As with the NBDs, the TMDs of Pdr5 are pseudo-dimeric (Fig. 1*D*, *Lower*). Each TMD contains five long α-helices that span the lipid bilayer, a broken transmembrane α-helix that makes a near-90° bend inside the membrane (TH5 in TMD1 and TH11 in TMD2), and an N-terminal amphipathic α-helix that lies on the membrane surface (TH1a and 7a) (Fig. 1D and Fig. S3). The latter two are also present in ABCG2, the closest structurally characterised relative of Pdr5 (*33*). The space between TMD1 and TMD2 is the location of the substrate channel in ABC transporters (*1, 34*). Two TMD α-helices are of particular importance to transport: the so-called coupling helix (CpH) and connecting helix (CnH) (*35*). In Pdr5, the CpHs are in the cytoplasmic portion of TH2 and TH8. The CnHs are the amphipathic helices TH1a and TH7a of TMD1 and TMD2 (Fig. 1*D* and Fig. S3). The helices form the main site of the interaction between the TMD and the NBD. In ABCG2, these are also pivot points for the structural transition between the substrate-bound (or nucleotide-free) and substrate-releasing (ATP-bound) conformations of the transporter (*33, 35*).

### Pdr5 contains a nucleotide-sensing linker domain

One distinctive feature of Pdr5 is the linker domain (LD), situated between the two halves of Pdr5 and composed of two distinct stretches. One part (LD1) is made from a loop extrusion of 30 amino acid residues situated between the first two β-sheet strands of NBD1; it also contains an amphipathic α-helix (H3) that rests on the surface of the lipid bilayer (Fig. 1*B-D*, Fig. S2 and Fig. S3). The other part (LD2) is formed by a polypeptide chain that links the C-terminus of TMD1 with the N-terminus of NBD2 and is shaped like an arch that rises from the surface of the bilayer, goes past the first stretch of the LD and ends with a short α-helical fragment (H17) that is almost perpendicular to the long helix in the domain. The 44 residues connecting the short helix H17 with the NBD2 could not be resolved, due to flexibility (Fig. 1*B-D*, Fig. S2 and Fig. S3). The base of the loop of the first stretch and the middle part of the arch in the second stretch directly contact the degenerated ATP-binding site in NBD1 (Fig. 1*B* and Fig. S4). In the apo-Pdr5, the arch of the LD2 connecting the TMD1 and NBD2 is not as well defined in the map, and the short helix H17 could not be resolved (Fig. S4). This could indicate that ATP binding in the degenerated site has a stabilising effect on that part of the LD, possibly through the contact of E804 with the ribose, as well as an electrostatic interaction of the Q801 side-chain with the triphosphate (at least in ADP-Pdr5) (Fig. S4 and Fig. S6). The unresolved region of Pdr5 between the H17 helix and NBD2 contains a ubiquitination site on K825 (*36, 37*), but there is no indication yet of other complex interactions in that part of the transporter.

Sequence alignment of the PDR family revealed that the linker region contacting ATP contains a motif, highly conserved in the PDR subfamily, with a consensus sequence MQKGEIL (Fig. S4) and the glutamine corresponding to Q801 mentioned above. The most highly conserved (95%) residue in the motif is the glutamate (E804 in Pdr5); in all other cases, this residue is an aspartate. In our structure, E804 contacts N1011 and, together with the two hydrophobic residues I805 and L806, positions the arch of LD2 between the loop extension of LD1 and NBD2 (Fig. S4). In the apo-Pdr5 structure, the residues of the conserved MQKGEIL motif cannot be discerned in the density, indicating that the motif plays a structural role, perhaps ensuring that the linker domain becomes more conformationally rigid when a nucleotide is bound in NBS1.

### Pdr5 contains a structurally unique extracellular domain

The extracellular (or apical) domain of Pdr5 (labelled as ECD1 and ECD2 on Fig. 1*B*) interrupts the TMDs just before their C-terminal α-helices: TH6 and TH12 (Fig. 1*B* and Fig. S3). Most of the ECD is composed of structured loops, with two short helices (ECD1 −H16 and ECD2-H29) at the base of the domain, and a slightly longer, distorted helix facing the extracellular milieu (ECD2-H28). Each half of the ECDs contains disulphide bridges (C722 and C742 in ECD1, and C1411 and C1455 as well as C1427 and C1452 of ECD2), which are probably involved in the stabilisation of the local fold. Interestingly, there is no interdomain disulphide bridge like that seen in ABCG2. In our cryo-EM map, the backbone of the asparagine N734 in ECD1 is well resolved, and its side-chain has additional density that is likely to be a site for glycosylation (Fig. S5), consistent with mass spectrometry profiling(*38*).

### NBS1 of Pdr5 does not support ATP hydrolysis

A number of important ABC transporters possess atypical NBS that have greatly diminished or no hydrolytic abilities due to substitutions of key residues in canonical motifs. In *Saccharomyces* sp. the three major drug efflux pumps (Yor1, Snq2, and Pdr5) have an atypical NBS alongside the canonical one. Endowing NBS1 of Pdr5 with hydrolytic capabilities abolishes substrate transport (*13*).

The non-catalytic site of Pdr5 differs from the canonical NBS2 in all but one of the conserved regions: the D-loop. The signature motif of the PDR family of ABC transporters, VSGGE, is present in NBD1, and correctly supplying the relevant amino acid residues to perform hydrolysis in NBS2, when the domains come together, as part of the catalytic cycle of ABC transporters. Vice versa, the signature motif of NBD2, LNVEQ, is mutated completely and does not complement NBS1 (Fig. 2*C*). However, perhaps the most consequential mutation is found in the Walker B motif of the deviant ATP-binding site. Where the canonical site has a glutamate residue (E1036), the deviant has an asparagine (N334) in the same position (Fig. 2*C*), with a tremendous impact on its ATPase activity. As the glutamate of Walker B polarises a water nucleophile in the catalytic site(*22*), the hydrolysis of ATP cannot be accelerated with the N334 substitution. In the canonical sites of full-length transporters, a similar inhibitory effect has been achieved by mutating the catalytic glutamate to glutamine, which resembles the protonated form of glutamate (formed upon polarising the water)(*39–41*). Notably, in our structures of Pdr5, the non-catalytic ATP-binding site does not contain magnesium (Fig. 2*C*).

**Fig. 2.**
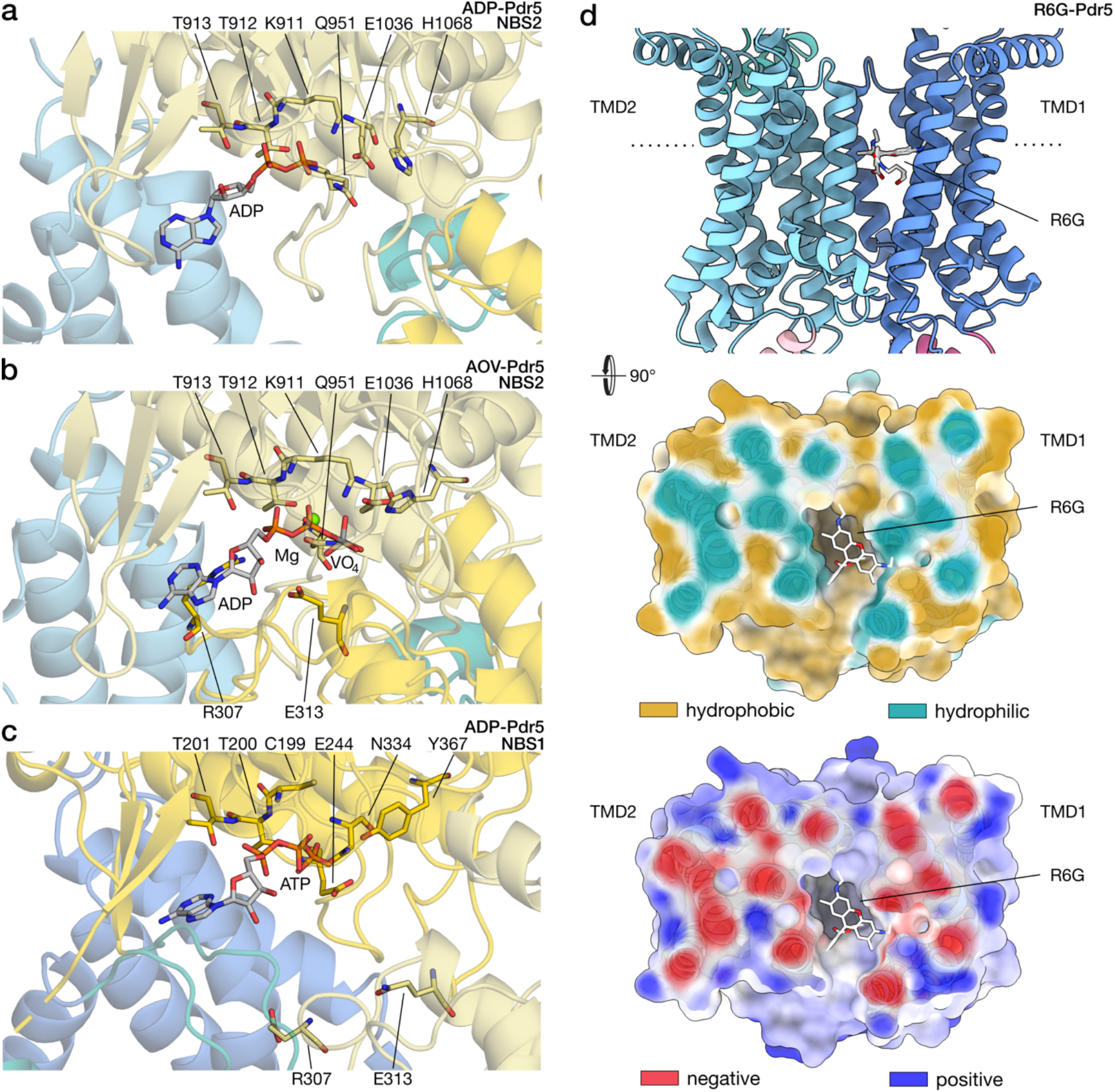
Nucleotide and substrate binding to Pdr5. The panels illustrate the contacts between Pdr5 with nucleotides (**A**-**C**) and rhodamine 6G (**D**). Some of the residues that are important for nucleotide binding and/or catalysis are highlighted and labelled (*cf*. Fig. S5). The domain colouring follows the scheme of the previous figure. (A) The catalytic site of Pdr5 in inward-facing conformation (ADP-Pdr5) with bound ADP. (B) The catalytic site of outward-facing Pdr5 (AOV-Pdr5) in a vanadate-trapped state that mimics the intermediate step of hydrolysis with an ADP-orthovanadate molecule. A co-ordinated magnesium ion is depicted as a green sphere. (C) The deviant, inactive site of Pdr5, showing the bound ATP. The residues surrounding the nucleotide are at the corresponding positions of the catalytic site. (D) Side view (*Top*) and cross-section (*Centre, Lower*) of the transmembrane part of Pdr5 in inward-facing conformation (Pdr5-R6G). Dotted line on the side-view denotes the position of the cross-section. The efflux substrate rhodamine 6G is bound in the entrance cavity between the TMD1 and TMD2 domains. The cross-section of Pdr5 is coloured by hydrophobicity, from turquoise (hydrophilic) to tan (most hydrophobic), and electrostatic potential, from red (negative) to blue (positive). Abbreviations: AOV, ADP-orthovanadate; R6G, rhodamine 6G; TMD, transmembrane domain.

### NBS2 of Pdr5 is a canonical catalytic site of an ABC transporter

The other nucleotide-binding site, NBS2, is a functional catalytic centre, capable of ATP hydrolysis and endowed with the canonical motifs that support enzymatic function(*5, 13, 22*), including the D-loop, which provides a crucial point of contact between the two halves of the NBD dimer, coupling ATP hydrolysis, NBD dimerisation, and substrate transport (*1, 42*).

Our structures of Pdr5 provide a glimpse into the catalytic site of the transporter in two forms: with bound ADP (ADP-Pdr5) and with an ADP-orthovanadate complex (AOV-Pdr5) (Fig. 2*A* and *B* respectively) in NBS2. The ADP-Pdr5 structure was solved using Pdr5 samples with added ATP, indicating that that the majority of ATP was hydrolysed before flash-freezing in liquid ethane. It also suggests that in the post-hydrolysis stage ADP remains bound to the catalytic site, making ADP-Pdr5 the default resting state of the transporter *in vivo*. We were able to mimic the hydrolysis transition state of the transporter using orthovanadate. The nucleotides bind the catalytic pocket in the manner similar to other ABC transporters, reflecting the high conservation of this site (*27, 33, 35*). Among residues that contact the nucleotide found in characteristic motifs, of particular importance are: K911, T912, and T913 in Walker A, with the lysine residue binding the terminal γ-phosphate of ATP; Q951 of the Q-loop, co-ordinating the magnesium ion in the ADP-V_i_ complex; E1036 of Walker B, mentioned before; H1068 of the H-loop, which acts as a proton-shuttle in the catalytic pocket; and residues R307 and E313 of the D-loop, which are part of NBD1 and bring together the two NBDs upon binding of ATP (Fig. 2*A* and *B*; Fig. S6). Pdr5 can utilise other nucleotides to fuel its transport activity (*14, 43*), and this can be rationalised by our structures from the distribution of nucleotide contacts.

### Pdr5 substrate rhodamine 6G binds in the cleft between the TMDs

We solved the molecular structure of Pdr5 with the ligand R6G, a common organic dye (*23*). R6G is nestled between the two TMDs, in the inward-facing conformation of Pdr5, with ADP in the catalytic NBS and ATP in the deviant site (Fig. *2D*). The amino-acid residues surrounding the ligand do not change positions considerably compared to the ADP-bound state (Fig. S6*C*). Because of this, we surmise that there is no cross-talk between the substrate binding pocket and the rest of the transporter in the inward-facing conformation, which is supported by the fact that Pdr5 is an uncoupled transporter (*22*).

R6G occupies a cleft that is positioned between the halves of the transmembrane portion of the transporter and is flanked by helices TMD1-TH2 and −TH5a as well as TMD2-TH8 and −TH11a (Fig. 2*D* *cf*. Fig. 1*D*). This includes residue S1360 which is known from mutational studies to be involved in R6G binding (*23*). The pocket makes a large opening that reaches more than halfway down the depth of the cell membrane and is accessible from the cytosol as well as the inner leaflet of the lipid bilayer (Fig. 3*A*). The surface of the pocket is a mixed hydrophobic/hydrophilic environment, with the predominance of the former (Fig. 2*D*). The studied transport substrates of Pdr5 include mostly lipophilic molecules, which seem compatible with the pocket, although it has been shown that substrate volume rather than lipophilicity is the major determinant of transporter selectivity (*7, 44*). The entrance to the substrate cavity on the side of the bilayer has a slight negative charge, opposite to the positively charged R6G, whilst the cavity itself remains more neutral electrostatically (Fig. 2*D*). The location of R6G in the cleft between the TMDs is comparable to the positioning of substrates in other multidrug transporters, such as ABCG2 (*45, 46*) or TmrAB (*27*). This suggests that this pocket, a feature of the inward-facing conformation of Pdr5, is the transporter’s substrate entry cavity (Fig. S7).

**Fig. 3.**
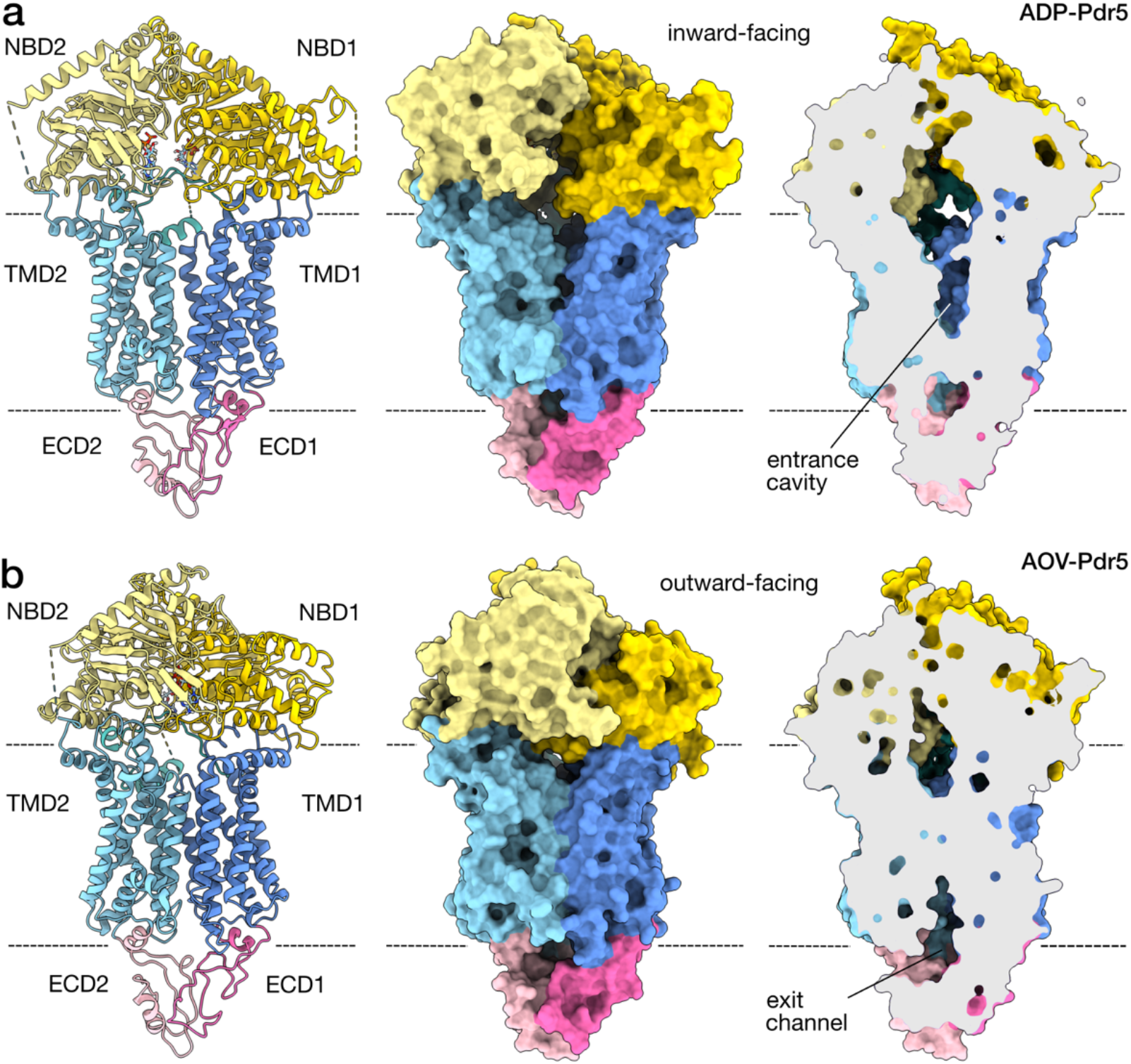
Conformational changes upon ATP binding. The two panels depict the molecular structure model of inward-facing (**A**) and outward-facing (**B**) conformations of Pdr5 in cartoon representation (*Left*), and as a solvent-excluded surface (*Centre*) with its cross-section (*Right*). (A) Shown is the Pdr5 in the post-hydrolytic state, with ADP present in the catalytic site (ADP-Pdr5) and the transport channel open onto the cytoplasmic side. The position of the substrate entrance cavity is indicated. (B) The Pdr5 in the vanadate-trapped, outward-facing state is depicted. The exit channel for the substrate, which opens onto the extracellular side, is indicated as well. Abbreviations: ECD, extracellular domain; NBD, nucleotide-binding domain; TMD, transmembrane domain.

### ATP binding induces a change to outward-facing conformation of Pdr5

The apo-Pdr5, ADP-Pdr5, and R6G-Pdr5 are in inward-facing conformations and are very similar (Fig. S8*A-B*); the differences are limited to small structural rearrangements around the nucleotide-binding sites when ADP and ATP are present. Our model of AOV-Pdr5, a product of vanadate-trapping of Pdr5, yielded a structural model that is in the outward-facing conformation (Fig. 3, Fig. S8*B* and Supplementary Movie 1). ADP and V_i_, when bound in the canonical NBS2, cause dimerization of the two NBDs. This occurs mainly through the interaction of the nucleotide, bound in NBS2, with the D-loop of the NBD1, and follows a pattern that is almost universally conserved amongst ABC transporters and other proteins that share a similar fold (*1, 11*). The interaction of the NBDs causes a structural change in the transmembrane and extracellular domains, switching the transporter’s overall conformation from inward-facing to outward-facing (Fig. 3).

In the outward-facing conformation, the helices of the TMD1 and TMD2 tilt towards each other on the cytosolic side and spread further apart on the extracellular side (Fig. 3*B*, *Left*). Helices that line the substrate entry cavity (TMD1 −TH2, −TH5a, and TMD2 −TH8, −TH11a) tighten around it, simultaneously opening up on the extracellular side to reveal a cleft that joins up with the newly expanded cavity between ECD1 and ECD2 (Fig. 3B, *Right*). The conformational state changes are highly asymmetric. NBD2, which contains the catalytic nucleotide-binding site, performs a greater domain movement than NBD1, which hosts the deviant site. Similarly, the greatest conformational changes in the transmembrane region involve the TMD2, which is responsible for the closure of the substrate cavity and the opening of the extracellular part of the transporter, while TMD1 is comparatively invariant (Fig. S8*B*). The single-site hydrolysis is a simplification of the symmetric ABC transporter system, whereby two ATP molecules are thought to complete the transport cycle (*47*).

The use of vanadate to mimic the transition state of nucleotide hydrolysis allows the transporter to be captured in a state that would otherwise be inaccessible due to the speed of hydrolysis (*24–27*). Various studies report a high degree of structural similarity of the transition and the pre-hydrolytic states for other ABC transporters, indicating that ATP binding and not hydrolysis induce the conformational change in the transporter from the inward-facing to the outward-facing (*27, 48–50*). The details of the substrate passage through the channel between the TMDs remain to be elucidated for Pdr5 (and indeed for other ABC transporters), but we envisage it to be somewhat akin to a peristaltic action where the closure of the entry channel on the cytoplasmic side pushes the substrate towards the expanding exit channel (Supplementary Movie 2). The access to both the entrance cavity and the exit channel is from the membrane-cytosol and membrane-extracellular interface (Fig. 3 and Fig. S7), which in principle can accommodate substrates coming from and/or directed to a lipophilic or hydrophobic environment; this appears compatible with the amphipathic character of reported Pdr5 substrates and implies that substrates are not released into the extracellular space but rather into the outer leaflet of the plasma membrane. The entrance and exit sites are asymmetric in shape and surface properties, so that the probability of entry from one site may be higher than from the other (Fig. 3). The Pdr5 exit channel might also contain the equivalent of the leucine gate found in ABCG2, which controls drug extrusion in that protein (*51*) (Fig. S9).

### ATP binding induces rearrangement in the N-terminal extension of Pdr5

The N-terminal part of Pdr5 contains a stretch of some 120 amino-acid residues that precedes the conserved portion of the NBD domain and is characteristic of the PDR subfamily. Mutagenesis data imply a role of this domain in signalling to other cellular processes and/or trafficking. We observe that the N-terminal extension contains two helices H1 and H2 connected by an unresolved region of 43 amino-acids (Fig. S2 and Fig. S3). The two helices are on the outside of NBD1 (top left of Fig. 1*C*; *cf*. Fig. S8*D*) and occupy well defined positions in the inward-facing state (ADP-Pdr5). In contrast to the rest of the NBD1, which maintains its overall structure between the two states, a pronounced structural change in the N-terminal extension is observed with ATP binding and switch to the outward-facing (AOV-Pdr5) conformation (Fig. S8*D*). In the outward-facing state, helix H1 appears to be disordered and is displaced by a coupled movement of H2 and helix H15 (Fig. S8*D*). These conformational changes could support recognition to trigger downstream effector processes.

### Drug efflux occurs as Pdr5 switches between inward- and outward-facing conformations

The structural models of Pdr5 in different nucleotide and ligand states presented here suggest how efflux works (summarised graphically in Fig. 4). In the resting state, Pdr5 adopts an inward-facing conformation with the substrate entry channel open towards the cytoplasm and inner leaflet of the membrane. In this state, corresponding to our ADP-Pdr5 model, the transporter has an ADP molecule in NBS2 and ATP in the deviant NBS1. On the basis of cellular concentrations of ATP in *S. cerevisiae* (1.51 ±0.32 mM (*52*)) and the transporter’s K_M_ for the nucleotide (1.7 to 1.9 mM *in vivo*^12,44^, 0.44±0.05 mM *in vitro* (*14*)), it seems likely that the apo state may represent about 50% of the population. The other Pdr5 states, which were prepared from samples with added ATP, have the nucleotide bound in the non-catalytic, deviant binding site; this would support earlier suggestions that ATP remains in the site throughout the transport cycle (*32*) under physiological conditions. These structures now also provide the basis to explain important functional mutants of Pdr5 (for a selection see Table S1).

**Fig. 4.**
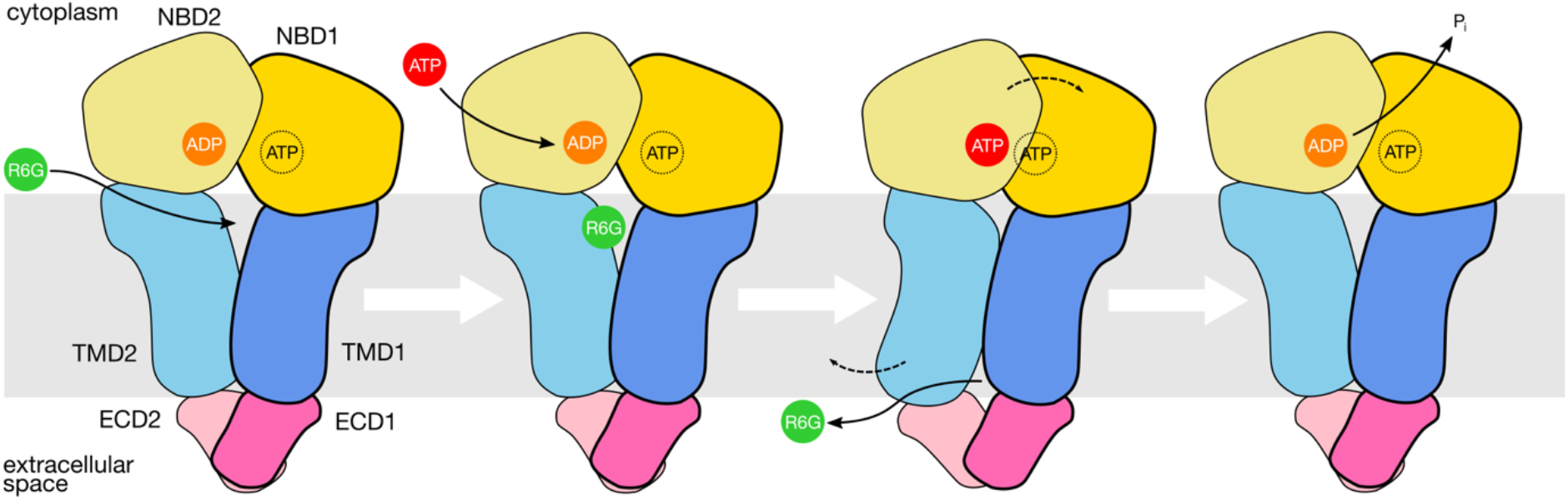
Transport cycle of S. cerevisiae Pdr5. The substrate (in this case, rhodamine 6G) enters the cavity between the transmembrane domains from the cytoplasmic side when Pdr5 is in inward-facing conformation. ADP remaining in the catalytic site from the previous hydrolysis step is exchanged for ATP. ATP binding triggers a conformation change in Pdr5 from inward-to outward-facing, whereby the substrate is pushed through the substrate channel and released into the extracellular medium. Upon nucleotide hydrolysis, the transporter reverts to inward-facing conformation, ready to receive another molecule of substrate. In cellular conditions, an ATP molecule remains bound to the inactive site throughout the cycle. The asymmetric nature of nucleotide hydrolysis means that one half of Pdr5 makes a larger conformational move than the other. Abbreviations: ECD, extracellular domain; NBD, nucleotide-binding domain; R6G, rhodamine 6G; TMD, transmembrane domain.

The substrate entry cavity of the inward-facing Pdr5 can receive the efflux substrate, as evidenced by our structure of Pdr5 with R6G (R6G-Pdr5). At this point, if ADP in the catalytic site is exchanged for ATP, transport can occur. The binding of ATP induces a structural change, which involves the transporter switching conformation from inward-facing to outward-facing, as exemplified by our vana-date-trapped structure (AOV-Pdr5). This change involves sealing off the substrate entry cavity and opening of an exit channel on the extracellular side of the transmembrane domain. The transport substrate moves from the entrance cavity into the exit channel and is released into the extracellular medium or outer leaflet. After the nucleotide is hydrolysed, the transporter reverts to the inward-facing conformation. In this state, ADP remains bound to the catalytic site, and the substrate entry channel is re-opened. Thus, the transport cycle of Pdr5 is completed, and the transporter can accept the next molecule of the substrate.

## Materials and Methods

### Protein Expression and Purification

The protocol of Wagner *et al*. (*14*) was used for producing and purifying stable Pdr5 preparations from *S. cerevisiae* and is summarised briefly here.

Histidine-tagged Pdr5 was purified from *S. cerevisiae* strain YRE1001 (*22*) grown in YPD medium at 30 °C. Cells were harvested by centrifugation at OD_600_ = 3.5 and lysed with glass beads in buffer containing 50 mM Tris-HCl (pH = 8.0), 5 mM EDTA and EDTA-free Protease Inhibitor Cocktail tablets (Roche AG, Basel, Switzerland). Cell debris was removed by centrifugation (4 °C): twice at 1,000 × g for 5 min and once at 3,000 × g. Cell membrane was pelleted from the resulting supernatant by centrifugation at 20,000 × g for 40 min (4 °C). The cell membrane pellet was resuspended in Buffer A: 50 mM Tris-HCl (pH = 7.8), 50 mM NaCl, 10% (w/v) glycerol and adjusted to 10 mg/mL total protein concentration. The solubilisation of membrane proteins was achieved through addition of 1% (w/v) of *trans*-4-(*trans*-4’-alkylcyclohexyl)cyclo-hexyl-α-D-maltoside (*trans*-PCC-α-M; Glycon Biochemicals, GmbH, Luckenwalde, Germany) and 1 h incubation at 4 °C, with gentle stirring. Non-solubilised material was removed by centrifugation at 170,000 × g for 45 min (4 °C).

Pdr5 was separated from other proteins by immobilised metal ion affinity chromatography (IMAC), using a HiTrap Chelating HP column (GE Healthcare, Chicago, IL, USA) loaded with Zn^2+^ ions and equilibrated with low-histidine buffer: 50 mM Tris-HCl (pH = 7.8), 500 mM NaCl, 10% (w/v) glycerol, 2.5 mM L-histidine, 0.003% w/v *trans*-PCC-α-M. The sample was loaded onto the column, washed, and eluted with a concentration gradient of histidine (up to 100 mM). Fractions of IMAC eluate containing Pdr5 were pooled and concentrated using a Vivaspin 6 centrifugal concentrator unit (Sartorius AG, Göttingen, Germany) with a molecular weight cut-off (MWCO) of 100 kDa. Concentrated protein was further purified using size-exclusion chromatography (SEC) on a Superdex 200 10/300 GL (GE Healthcare, Chicago, IL, USA) column containing 0.003% (w/v) *trans*-PCC-α-M.

Fractions of highest Pdr5 concentration (approx. 2.5 mg/mL) from SEC were selected for reconstitution into peptidiscs (*17*). The peptidisc solution was prepared from bulk lyophilised peptidisc (Peptidisc Biotech, Vancouver, BC, Canada) dissolved in 20 mM Tris-HCl (pH = 8.0) to a final concentration of 10 mg/mL. Dissolved peptidisc was mixed with solubilised Pdr5 sample in 1:1 weight ratio (1:38 molar ratio) and incubated for 30 min at room temperature. The mixture was separated using SEC on a Superose 6 10/300 GL column (GE Healthcare, Chicago, IL, USA) equilibrated with buffer containing 50 mM Tris-HCl (pH = 7.8), 100 mM NaCl and no detergent. This step yielded homogenous Pdr5/peptidisc assemblies.

### Cryo-EM Sample Preparation and Data Collection

To prepare the vanadate-trapped state sample (AOV-Pdr5), Pdr5 resuspended in peptidiscs was mixed with 2 mM of ATP and 2 mM MgCl_2_, and incubated for 2 min. To this solution, 200 μM of freshly prepared sodium orthovanadate (Na_3_VO_4_) was added and incubated for further 2 min. In the case of the R6G-Pdr5 dataset, Pdr5 was mixed with 2 mM of ATP, 2 mM MgCl_2_, and 100 μM of rhodamine 6G solution in water. In the case of the ADP-Pdr5, just 2 mM of ATP, 2 mM MgCl_2_ were added. No additives were used to prepare the apo state sample. To improve the distribution of particles in vitreous ice, an aqueous solution of CHAPSO was added to all the samples to final concentration 8 mM. The solution was mixed by pipetting and used immediately in grid preparation. 3 μL of the sample mixture were applied onto a Cu 300-mesh EM grid with R1.2/1.3 holey carbon support film (Quantifoil Micro Tools GmbH, Großlöbichau, Germany) which had been glow-discharged prior to use. The sample applied to the grid was vitrified in liquid ethane at −180°C using a Vitrobot Mark IV (Thermo Fischer Scientific Corp., Eindhoven, The Netherlands).

The grids were imaged using Titan Krios G3 TEM (Thermo Fischer Scientific Corp., Eindhoven, The Netherlands) operated at liquid nitrogen temperature, 300-kV accelerating voltage, and in EF-TEM mode with GIF slit size of 20 μm. The micrographs were recorded using automated image acquisition software EPU (Thermo Fischer Scientific Corp., Eindhoven, The Netherlands), in the case of the ADP-Pdr5, R6G-Pdr5, and AOV-Pdr5 samples, on a K3 direct electron detector (Gatan Inc., Pleasanton, CA, USA) in super-resolution counting mode, at a detector pixel size of 0.326 Å (nominal magnification 130,000×) and nominal defocus values between −0.7 and −2.2 μm. The micrographs of the apo-Pdr5 sample were collected with a K2 Summit direct electron detector (Gatan Inc., Pleasanton, CA, USA) in counting mode, at a detector pixel size of 1.07 Å (nominal magnification 130,000×) at similar nominal defocus values. For the apo-Pdr5 sample, exposures lasted 12 s and were collected in 40 fractions (processing frames) each, with total measured electron fluency of exposure was 58.1 e^-^/Å^2^. A total of 4,211 micrographs were acquired over the course of a single 72-hour data collection session. For the ADP-Pdr5 sample, the exposures lasted 1.31 s, collected in 49 fractions and fluency of 47.21 e^-^/Å^2^. This dataset contained 3,415 micrographs. The R6G-Pdr5 and AOV-Pdr5 datasets were obtained using similar exposure conditions, with fluencies of 47.73 and 48.13 e^-^/Å^2^; they amounted to 3,604 and 8,370 micrographs respectively.

### Cryo-EM Data Processing for Structure Solution

The cryo-EM map of apo-Pdr5 was reconstructed in RELION-3.0 (*28*) and cryoSPARC-2.15 (*53*). Micrographs were corrected for beam-induced sample motion within RELION. Contrast transfer function (CTF) parameters were estimated using Gctf-1.08 (*54*) on non-dose-weighted micrograph averages. A subset of the images with estimated maximum resolution below 4 Å was selected from the full dataset for further processing. A manual pick of ca. 2,000 particles was used to prepare a set of 2D references for automated particle search, which was subsequently optimised to yield diverse projections of Pdr5 particles. The initial particle pool was extracted in 320-pixel boxes and pruned through a series of reference-free 2D classification rounds to 68,154. A model for cryo-EM reconstruction was created through stochastic gradient descent (SGD) algorithm in RELION on 8,000 randomly selected particles. The model and the pruned particle pool were subjected to 3D auto-refinement, which generated the first intermediate cryo-EM map used for subsequent Bayesian particle polishing and CTF refinement (*55*).

To improve the quality of the protein part of the apo-Pdr5/peptidisc assembly cryo-EM map, the density of the peptidisc was subtracted from the full map, using tools and procedures implemented in RELION (*56, 57*). First, a mask template was prepared by removing the Pdr5 part of the density from the full map, with the Volume Eraser utility of UCSF Chimera-1.13 (*58*). Second, the template was transformed into a binarised mask with a cosine-soft edge in RELION. The signal from within the mask was subtracted from the particles in the full map set from the last 3D autorefinement step. The result of the subtraction was validated with 2D classification of the subtracted particle set. The final map reconstruction was performed on subtracted particles using non-uniform refinement (NU-refinement) algorithm in cryoSPARC (*59*). The overall resolution of the obtained map, as judged by Fourier-shell correlation (FSC) of the half-maps with the gold-standard FSC_0.143_ criterion (*29*), was 3.45 Å.

The cryo-EM maps from the ADP-Pdr5 and the rhodamine 6G R6G-Pdr5 datasets were processed similarly, although RELION-3.1 was used for the reconstruction and corrected for high-order aberrations and anisotropic magnification errors (*60*). The raw micrographs were down-sampled by a factor of 2, yielding an input set of images with an effective pixel size of 0.652 Å. Contrast transfer function (CTF) parameters were estimated using CTFFIND-4.1 (*61*) on non-dose-weighted micrograph averages. The initial particle pool was extracted in 524-pixel boxes and pruned through a series of 2D and 3D classification rounds. In the case of the ADP-Pdr5 dataset, an improvement in the signal-to-noise ratio of the protein part of the Pdr5/peptidisc assembly was achieved using SIDESPLITTER (*62*) which locally de-noises cryo-EM maps (*63*) and 3D map reconstructions were done externally to RELION-3.1 between the iterative steps of 3D auto-refinement. The overall resolution of the final maps obtained by the combined use of SIDESPLITTER and RELION-3.1 was 2.85 Å (ADP-Pdr5) and 3.13 Å (R6G-Pdr5), per the FSC_0.143_ criterion.

The vanadate-trapped AOV-Pdr5 dataset was processed similarly to apo-Pdr5, combining the use of RELION-3.1 and cryoSPARC-3.0. The raw micrographs were down-sampled to an effective pixel size 0.652 Å. The initial particle pool was extracted in 524-pixel boxes and pruned through a series of 2D and 3D classification rounds. The overall resolution of the final map was 3.75 Å, per the FSC_0.143_ criterion.

### Model Building and Refinement

The reconstructed cryo-EM map of ADP-Pdr5 was used for *de novo* building of an atomic model of *S. cerevisiae* Pdr5. A preliminary model was calculated using the Rosetta software suite (*64*).The model was adjusted manually in Coot-0.9 (*65*), part of the CCP-EM package (*66*), and in ISOLDE (*67*). Model refinement was carried out in real space using PHENIX-1.18.2 (*68*). To assist model building, the maps were subjected to a density modification procedure (*69*), implemented as part of the PHENIX program suite. Model building in high flexibility regions of Pdr5 was facilitated through local map sharpening with LocScale (*70*), part of the CCP-EM program suite (*66*). Other models were built manually, using the ADP-Pdr5 structure as a starting point. Locally sharpened maps were also used for visualisation purposes. Molecular graphics were rendered in ChimeraX-1.0 (*71*) and PyMol 1.8.2.3. All refinement and data collection statistics are summarised in Table S2.

## Acknowledgements

We thank the staff of the Bio-cEM facility at the Department of Biochemistry, University of Cambridge, Dr Dimitri Y. Chirgadze, Dr Steven Hardwick, and Mr Lee Cooper for assistance with cryo-EM data collection.

## Author contributions

Conceptualization: BFL, LS

Methodology: AH, MW, DD, BFL, LS

Investigation: AH, MW, DD, SR, BFL, LS

Visualization: AH, DD, BFL, LS

Funding acquisition: HG, SHJS, BFL, LS

Project administration: BFL, LS

Supervision: AH, SHJS, BFL, LS

Writing – original draft: AH, BFL, LS

Writing - review & editing: AH, MW, DD, SR, HG, SHJS, BFL, LS

## Author contributions

Authors declare that they have no competing interests.

## Data and materials availibility

All molecular structures have been deposited in the PDB database under the following accession codes (PDB-IDs): XXXX, XXXX, XXXX, XXXX. Cryo-EM maps are available via EMDB: EMD-XXXXX, EMD-XXXXX, EMD-XXXXX, EMD-XXXXX

---

**Fig. S1.**
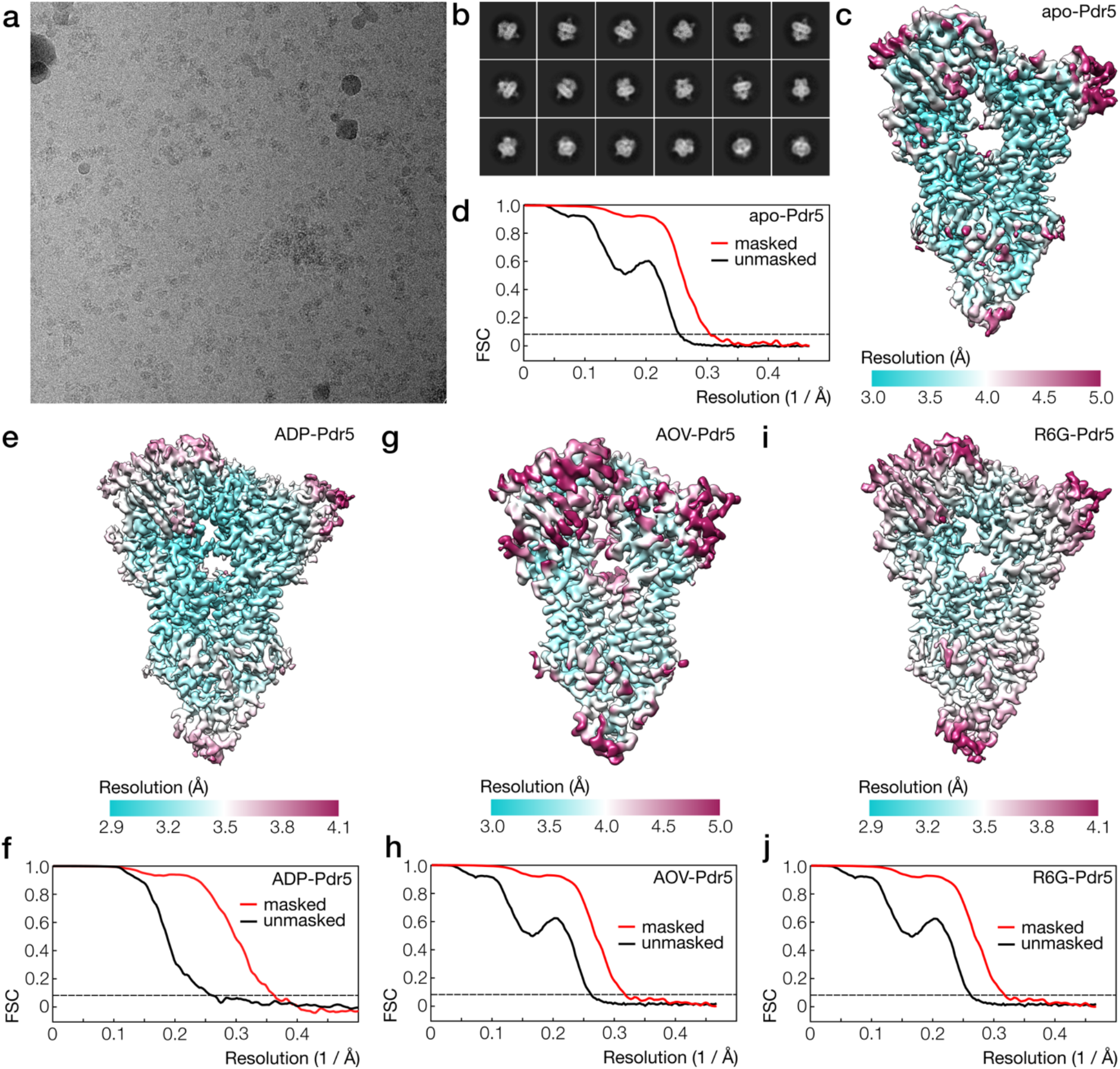
Cryo-EM imaging of Pdr5. Figure panels illustrate some of the steps in the structural reconstruction of apo-Pdr5 (**A-D**), ADP-Pdr5 (**E-F)**, AOV-Pdr5 (**G-H**), and R6G-Pdr5 (**I-J**). (A) Cryo-EM micrographs of Pdr5/peptidisc particles in vitrified ice. (B) Averaged images of the particles, obtained in reference-free 2D classification. (C) Cryo-EM map of apo-Pdr5, coloured by local resolution (see key). (D) Gold-standard FSC curves showing the correlation between the half-datasets in the unmasked (black line) and masked (red line) maps. The overall resolution was judged using the FSC_0.143_ threshold (dashed line). (E-J) Cryo-EM maps and FSC curves for the other Pdr5 reconstruction, analogous to (C–D). See also: Table S2. Abbreviations: FSC, Fourier-shell correlation.

**Fig. S2.**
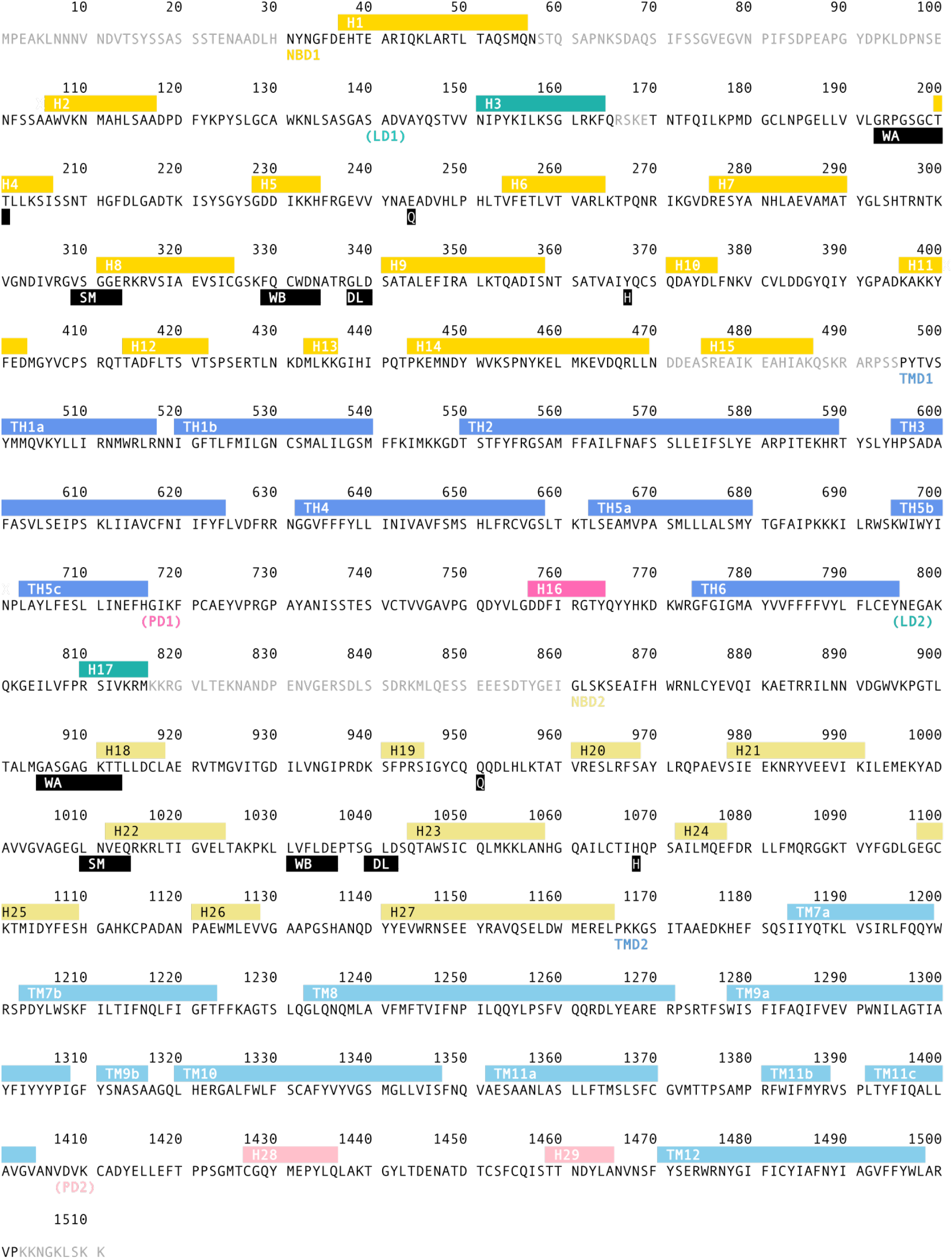
Annotated amino-acid sequence of Pdr5. Indicated are conserved features of the nucleotide binding sites (black boxes) and positions of α-helices (coloured boxes). The domains of Pdr5 are indicated, and the helices are coloured according to the scheme used elsewhere in the paper. Grey residues indicate parts of the ADP-Pdr5 model that could not be resolved in the cryo-EM map. Abbreviations: DL, D-loop; ECD, extracellular domain; H, H-loop; LD, linker domain; NBD, nucleotide-binding domain; Q, Q-loop; TMD, transmembrane domain; WA, Walker A (or P-loop); WB, Walker B (or A-loop).

**Fig. S3.**
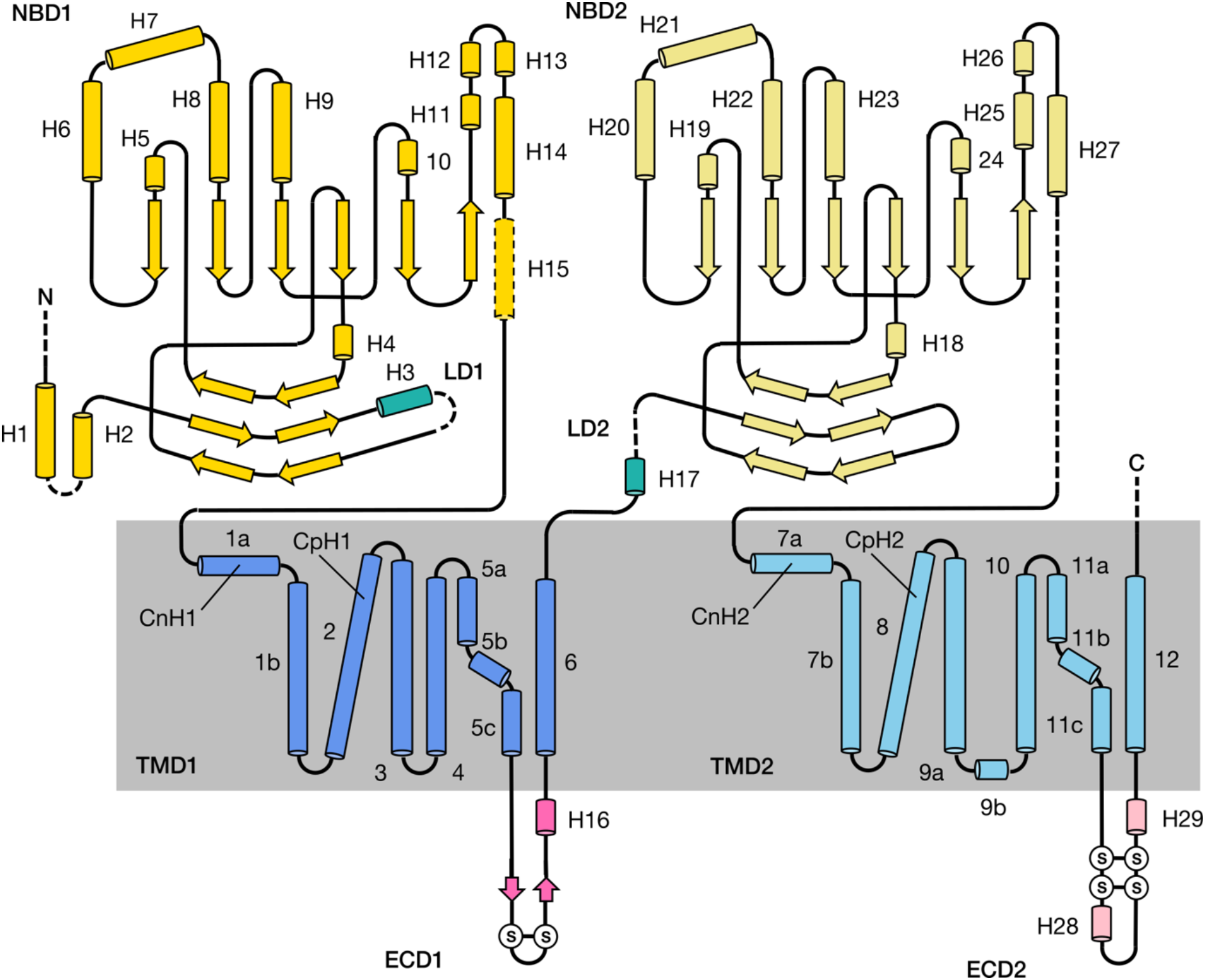
Domain topology of Pdr5. A schematic representation of domain architecture of Pdr5 showing the arrangement of its secondary structural elements. Cylinders and arrows represent alpha-helices and beta-sheets. The elements on the diagram are not shown to scale, but their relative positioning in the two halves of the protein reflects the evolutionary relation of the transporter domains through a geneduplication event. The colour scheme for domains is similar to the one used elsewhere in text. The dashed lines represent structure fragments that could not be resolved in the ADP-Pdr5 map (*cf*. Fig. S2). The circled letters “S” denote the position of disulphide bridges. According to the alternative nomenclature proposed for the PDR family (*3*), ECD1 and ECD2 are respectively EC3 and EC6. Abbreviations: ECD, extracellular domain; LD, linker domain; NBD, nucleotide-binding domain; TMD, transmembrane domain.

**Fig. S4.**
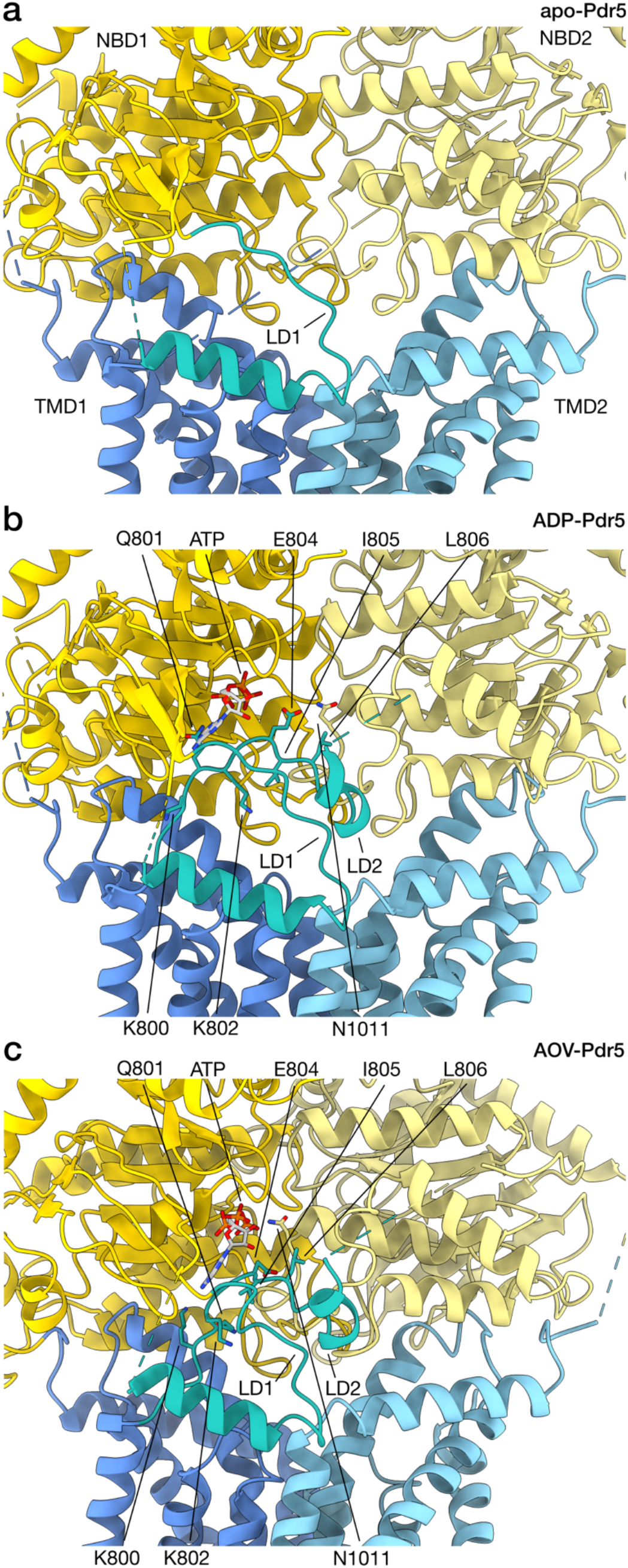
Structure of Pdr5 linker domain. Section of the Pdr5 structure surrounding the linker domain is shown in cartoon representation. The ATP in the non-hydrolytic site and some of the conserved residues of the linker domain are depicted in stick representation. The three panels represent different states of Pdr5: (**A**) the inward-facing apo-Pdr5, (**B**) the inward-facing ADP-Pdr5, (**C**) the outward-facing AOV-Pdr5. Abbreviations: LD, linker domain; NBD, nucleotide-binding domain; TMD, transmembrane domain.

**Fig. S5.**
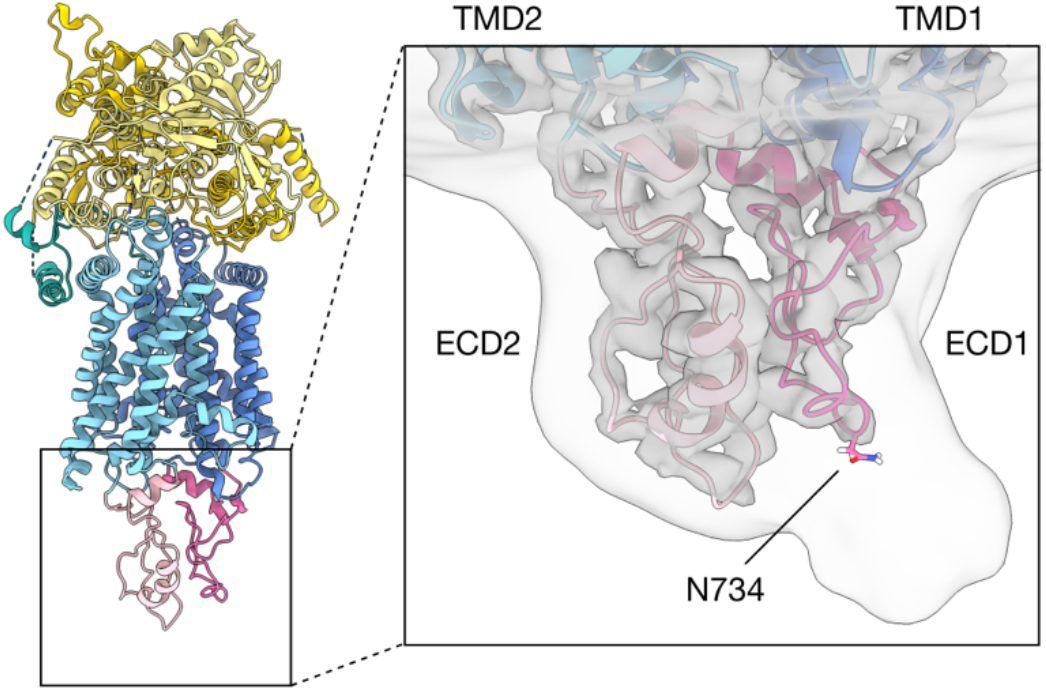
Glycosylation of Pdr5. Figure shows the side view of the Pdr5 (*Left*) and, in inset (*Right*), a putative glycosylation site on the extracellular domain of Pdr5. The two transparent surface representations of the apical portion of Pdr5 are from the same ADP-Pdr5 cryo-EM map, contoured at two different levels. The inner surface shows the Coulomb potential map immediately surrounding the protein; the outer is a noisier and Gaussian-filtered (2σ) version of the map, showing the protrusion around the asparagine residue (labelled), previously suggested to be glycosylated. Abbreviations: ECD, extracellular domain; TMD, transmembrane domain.

**Fig. S6.**
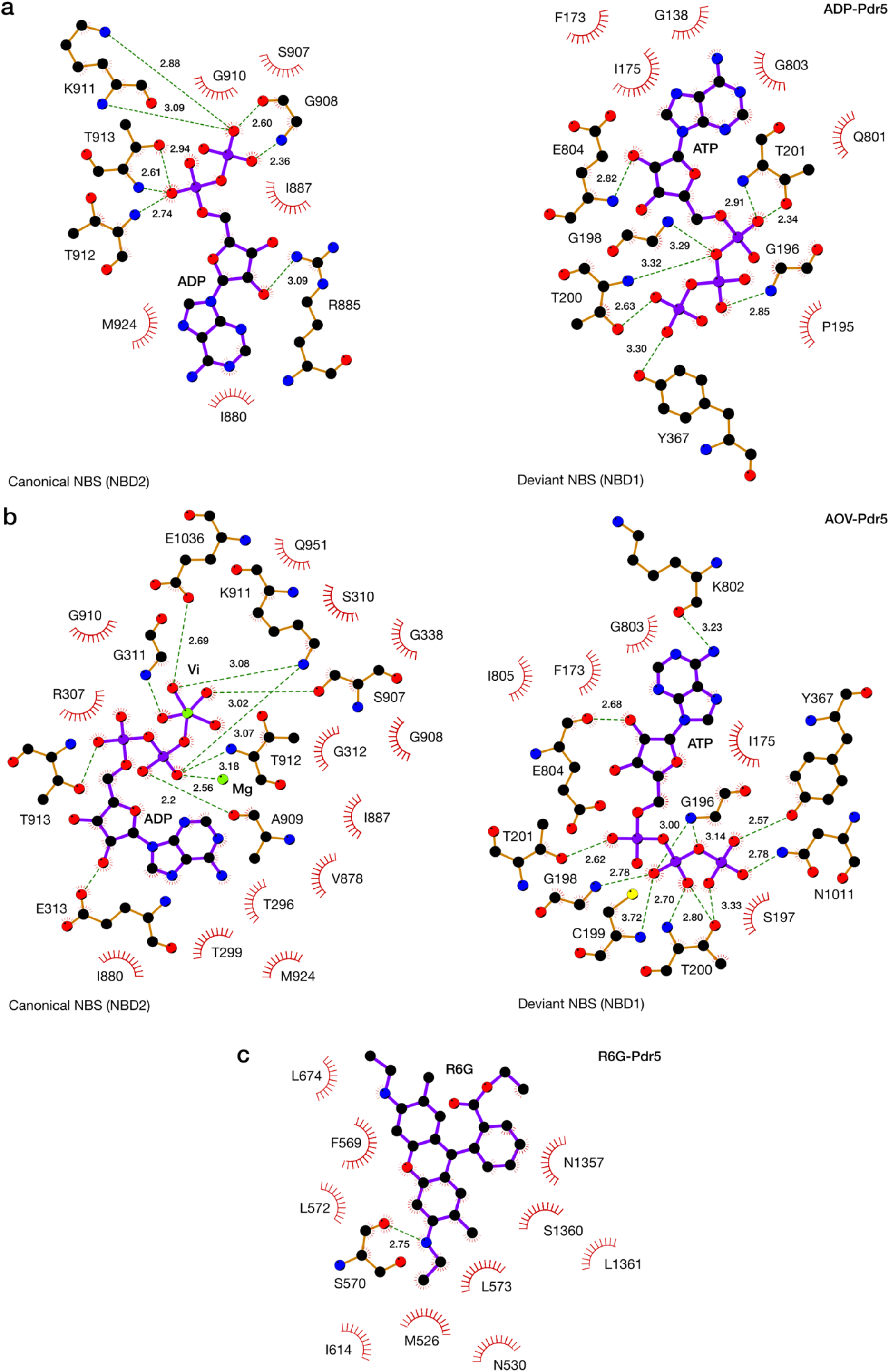
Details of protein-ligand interactions in Pdr5. Schematic representation of the network of molecular interactions between the nucleotides and the protein in the canonical and inactive nucleotide-binding sites of the inward-facing ADP-Pdr5 (**A**) and the outward-facing AOV-Pdr5 (**B**). The interactions between the transport substrate rhodamine 6G and the residues in the substrate entry channel in the inward-facing R6G-Pdr5 are also shown (**C**). The diagrams have been prepared with LigPlot+ (*72*). Atoms are depicted as coloured balls (C, black; N, blue; O, red; P; purple; S, yellow; Mg, V, green), ligand covalent bonds are purple and protein bonds are brown. Dashed green lines represent hydrogen bonds or other electrostatic interactions, with inter-atom distances indicated in Å. The radiating arc symbol represents amino-acid residues involved in hydrophobic contacts. Abbreviations: R6G, rhodamine 6G.

**Fig. S7.**
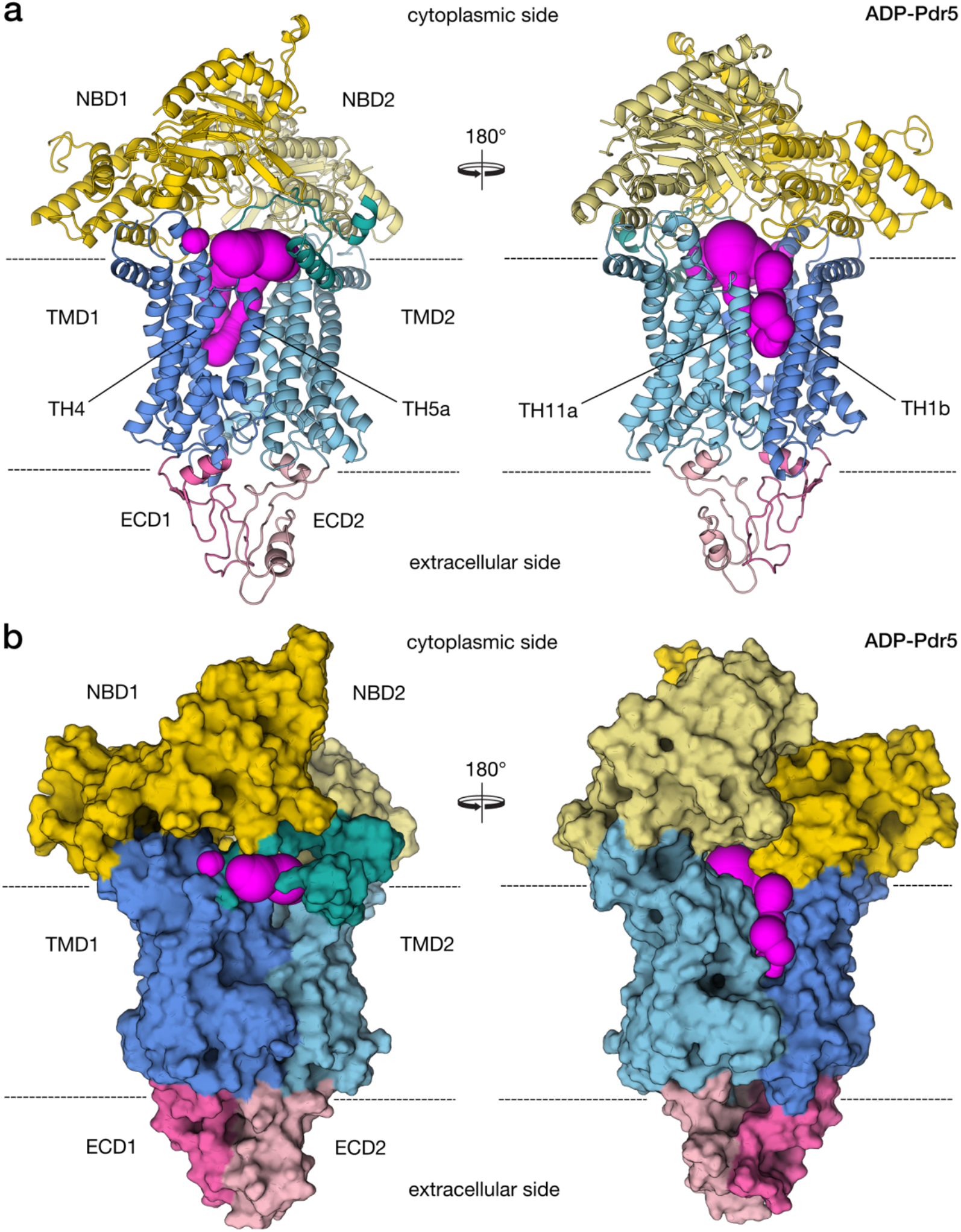
Analysis of potential entry sites in the inward-facing conformation of Pdr5. (**A**) Cartoon representation of the inward-facing state and the present tunnels (magenta) as determined by CAVER-3.0 (*73*). (**B**) Surface representation of Pdr5. This analysis demonstrates that Pdr5 has only one membrane entry site between helices TH1b and TH11a (A and B, *Right*), while the corresponding entrance on the opposite side (A and B, *Left*) is closed by TH4 and TH5a. The position of the membrane is indicated by dashed black lines.

**Fig. S8.**
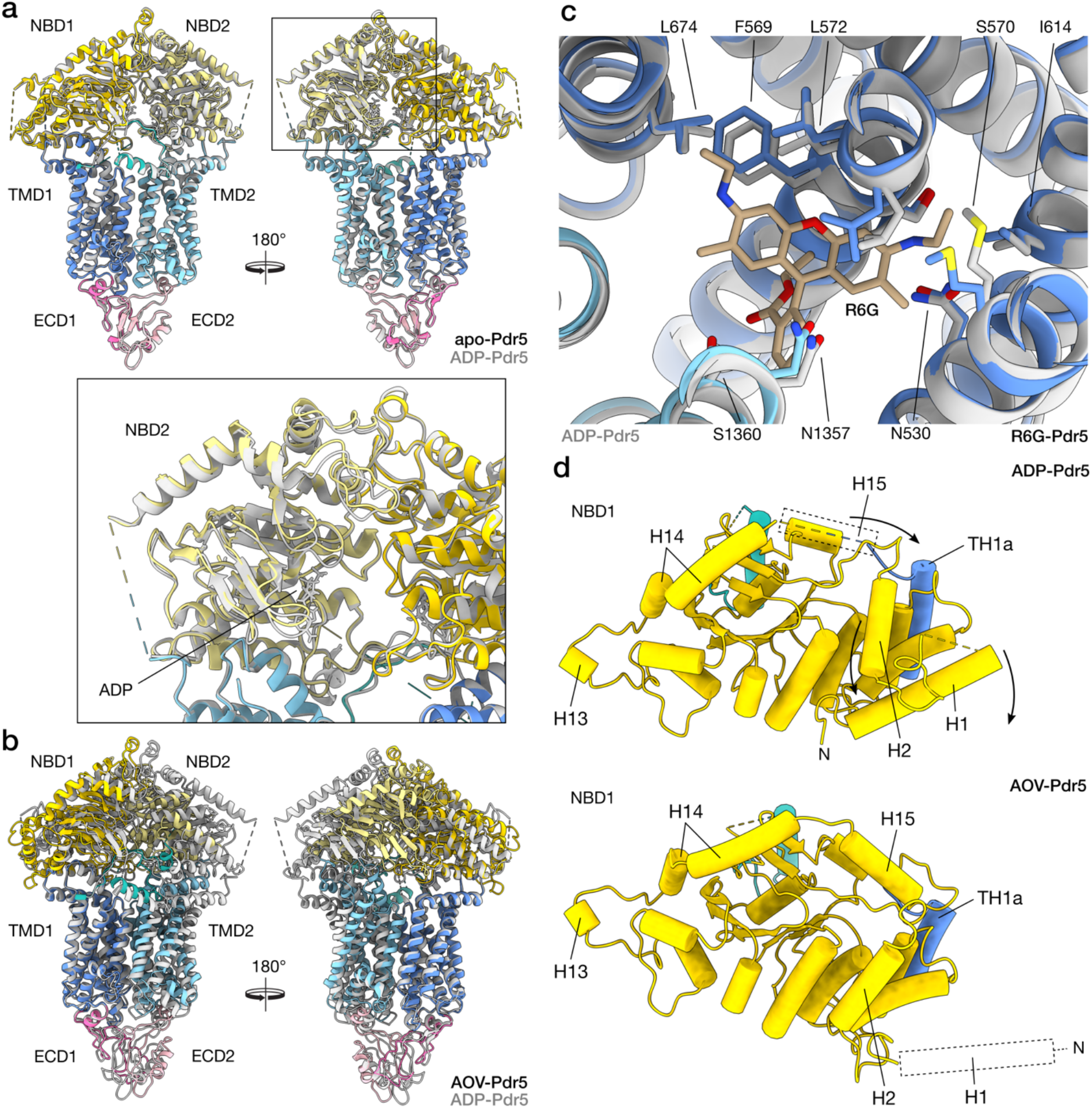
Structural differences between Pdr5 models. Shown here are structural overlays of Pdr5 models in cartoon representation: (**A**) between apo-Pdr5 and ADP-Pdr5, with the inset showing details of the NBD2 domain around the catalytic NBS; (**B**) AOV-Pdr5 and ADP-5 Pdr5 (or outward-facing and inward-facing conformations); (**C**) between R6G-Pdr5 and ADP-Pdr5 in the region surrounding the rhodamine 6G transport substrate. In panels A–C, ADP-Pdr5 structure is used as a reference and is coloured grey. Other models follow the domain colouring scheme used elsewhere in the paper. (**D**) The diagram depicts the difference between the conformation of NBD1 domain of Pdr5 in outward- and inward-facing 10 models (ADP-Pdr5 and AOV-Pdr5). In ADP-Pdr5, the N-terminal helices H1 and H2 are clearly resolved in the cryo-EM map, unlike helix H15, which did not adopt a discrete conformation. In AOV-Pdr5, H15 moves towards transmembrane helix TH1a, displacing H2 and H1, the latter of which is no longer visible in the map. Dashed lines denote the approximate position of unresolved helices, and arrows are an indication of movement. Abbreviations: ECD, extracellular domain; NBD, nucleotide-binding domain; R6G, rhodamine 6G; TMD, transmembrane domain.

**Fig. S9.**
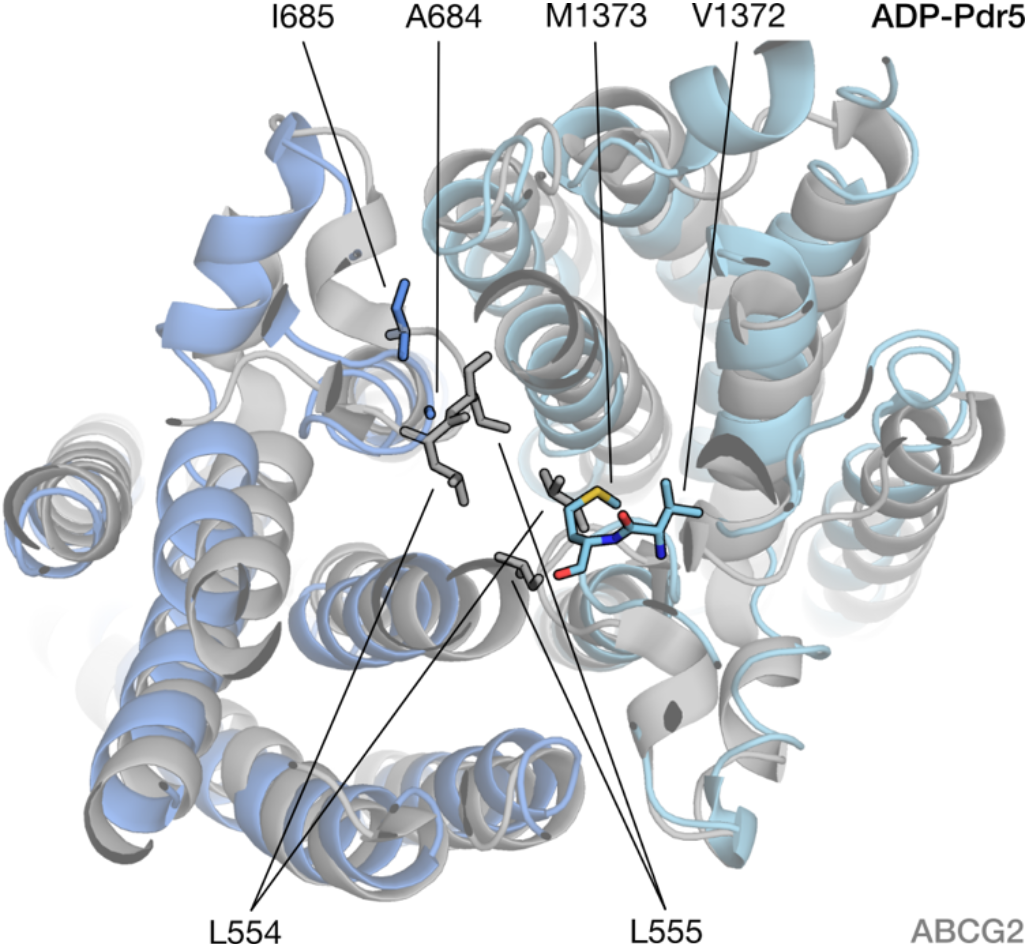
Leucine gate of human ABCG2 and the corresponding region of Pdr5. Human ABCG2 (grey, PDB-ID: 6hco) and Pdr5 (coloured) were superimposed and the residues of ABCG2 forming the di-leucine gate (L554 and L555) are highlighted (ABCG is a homodimer). The corresponding residues of Pdr5 (A684/I685 and V1372/M1373) are also highlighted.

**Table S1.** Selected mutations of Pdr5 with functional impact and possible structural explanation of the observed functional modulation of Pdr5.

| **Mutation** | **Location** | **Modulation of Pdr5 function** | **Reference** | **Explanation based on the structures** |
| --- | --- | --- | --- | --- |
| **E244G** | Q-loop of NBD1 | Increased drug sensitivity | (74) | Interaction with D-loop of NBD1 |
| **T257I** | H6 | Alters drug specificity | Uniprot P33302 | Not evident |
| **G302D** | Loop preceding the C-loop of NBD1 (X-loop) | Generalized drug resistance | Uniprot P33302 | Stabilizes the conformation of the C-loop of NBD1 |
| **S558Y** | TH2 | Impaired transport activity | (74) | Interacts with the linker TM11a-TM11b |
| **S648F** | TH4 | Alters drug specificity | Uniprot P33302 | Forms a helical bundle with TM5c and TM6 |
| **G905S** | Walker A of NBD2 | Inactivates transport | Uniprot P33302 | ATP binding / hydrolysis |
| **G908S** | Walker A of NBD2 | Inactivates transport | Uniprot P33302 | ATP binding / hydrolysis |
| **Q951Q** | Q-loop of NBD2 | Increased drug sensitivity | (75) | ATP binding / hydrolysis |
| **G1009C** | Loop preceding the C-loop of NBD2 (X-loop) | Generalized drug resistance | Uniprot P33302 | Interaction with LD1 (QKGEIL) |
| **G1040D** | D-loop of NBD2 | Alters drug specificity | Uniprot P33302 | Impaired NBD-NBD communication |
| **D1042N** | D-loop of NBD2 | Alters drug specificity | (32) | Impaired NBD-NBD communication |
| **S1048V** | H23 | Alters drug specificity | Uniprot P33302 | Not evident |
| **Y1311S** | TM9b | Alters drug specificity | Uniprot P33302 | Cluster of Y residues (1301, 1305, 1306, 1311) |
| **S1360F** | TM11a | Alters drug specificity | (76, 77) | Part of the R6G binding site |
| **S1386A** | TM11a | Creates bi-directional transport | (78) | On top of the R6G binding site |
| **T1393I** | TM11c | Alters drug specificity | Uniprot P33302 | On top of the R6G binding site |

**Note:** The multiple suppressor mutants of Pdr5, which have been described in the literature (see, for example<sup>2,7</sup>), have been excluded due to potential additive effects that cannot be predicted from the wild-type structure.

**Table S2.**
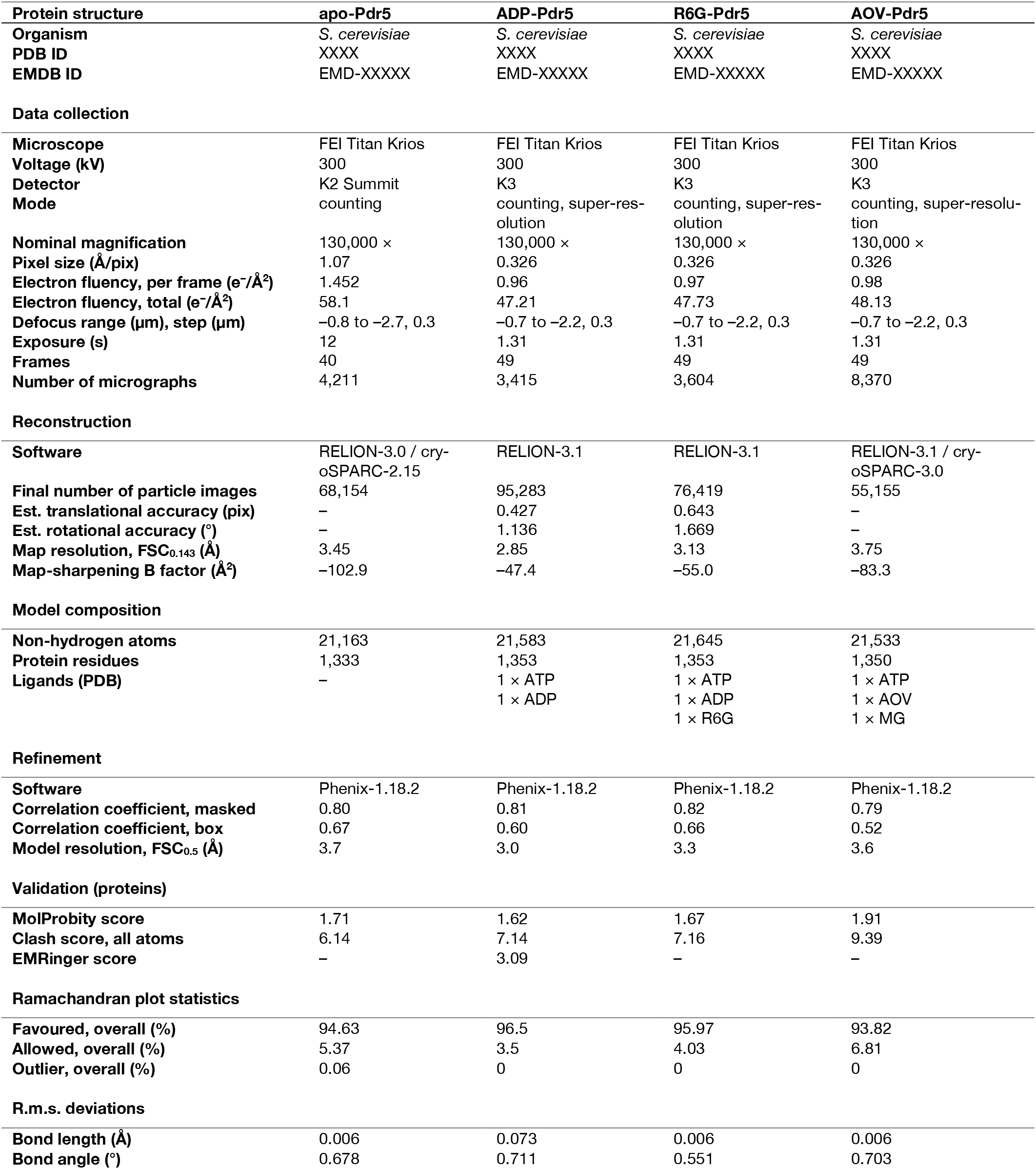
Cryo-EM data collection and refinement statistics

| Protein structure | apo-Pdr5 | ADP-Pdr5 | R6G-Pdr5 | AOV-Pdr5 |
| --- | --- | --- | --- | --- |
| Organism | *S. cerevisiae* | *S. cerevisiae* | *S. cerevisiae* | *S. cerevisiae* |
| PDB ID | XXXX | XXXX | XXXX | XXXX |
| EMDB ID | EMD-XXXXX | EMD-XXXXX | EMD-XXXXX | EMD-XXXXX |
| **Data collection** |  |  |  |  |
| Microscope | FEI Titan Krios | FEI Titan Krios | FEI Titan Krios | FEI Titan Krios |
| Voltage (kV) | 300 | 300 | 300 | 300 |
| Detector | K2 Summit | K3 | K3 | K3 |
| Mode | counting | counting, super-resolution | counting, super-resolution | counting, super-resolution |
| Nominal magnification | 130,000 × | 130,000 × | 130,000 × | 130,000 × |
| Pixel size (Å/pix) | 1.07 | 0.326 | 0.326 | 0.326 |
| Electron fluency, per frame (e<sup>-</sup>/Å<sup>2</sup>) | 1.452 | 0.96 | 0.97 | 0.98 |
| Electron fluency, total (e<sup>-</sup>/Å<sup>2</sup>) | 58.1 | 47.21 | 47.73 | 48.13 |
| Defocus range (µm), step (µm) | -0.8 to -2.7, 0.3 | -0.7 to -2.2, 0.3 | -0.7 to -2.2, 0.3 | -0.7 to -2.2, 0.3 |
| Exposure (s) | 12 | 1.31 | 1.31 | 1.31 |
| Frames | 40 | 49 | 49 | 49 |
| Number of micrographs | 4,211 | 3,415 | 3,604 | 8,370 |
| **Reconstruction** |  |  |  |  |
| Software | RELION-3.0 / cryoSPARC-2.15 | RELION-3.1 | RELION-3.1 | RELION-3.1 / cryoSPARC-3.0 |
| Final number of particle images | 68,154 | 95,283 | 76,419 | 55,155 |
| Est. translational accuracy (pix) | – | 0.427 | 0.643 | – |
| Est. rotational accuracy (°) | – | 1.136 | 1.669 | – |
| Map resolution, FSC<sub>0.143</sub> (Å) | 3.45 | 2.85 | 3.13 | 3.75 |
| Map-sharpening B factor (Å<sup>2</sup>) | -102.9 | -47.4 | -55.0 | -83.3 |
| **Model composition** |  |  |  |  |
| Non-hydrogen atoms | 21,163 | 21,583 | 21,645 | 21,533 |
| Protein residues | 1,333 | 1,353 | 1,353 | 1,350 |
| Ligands (PDB) | – | 1 × ATP<br>1 × ADP | 1 × ATP<br>1 × ADP<br>1 × R6G | 1 × ATP<br>1 × AOV<br>1 × MG |
| **Refinement** |  |  |  |  |
| Software | Phenix-1.18.2 | Phenix-1.18.2 | Phenix-1.18.2 | Phenix-1.18.2 |
| Correlation coefficient, masked | 0.80 | 0.81 | 0.82 | 0.79 |
| Correlation coefficient, box | 0.67 | 0.60 | 0.66 | 0.52 |
| Model resolution, FSC<sub>0.5</sub> (Å) | 3.7 | 3.0 | 3.3 | 3.6 |
| **Validation (proteins)** |  |  |  |  |
| MolProbity score | 1.71 | 1.62 | 1.67 | 1.91 |
| Clash score, all atoms | 6.14 | 7.14 | 7.16 | 9.39 |
| EMRinger score | – | 3.09 | – | – |
| **Ramachandran plot statistics** |  |  |  |  |
| Favoured, overall (%) | 94.63 | 96.5 | 95.97 | 93.82 |
| Allowed, overall (%) | 5.37 | 3.5 | 4.03 | 6.81 |
| Outlier, overall (%) | 0.06 | 0 | 0 | 0 |
| **R.m.s. deviations** |  |  |  |  |
| Bond length (Å) | 0.006 | 0.073 | 0.006 | 0.006 |
| Bond angle (°) | 0.678 | 0.711 | 0.551 | 0.703 |

## References and Notes

1. K. P. Locher, Mechanistic diversity in ATP-binding cassette (ABC) transporters. Nature Structural & Molecular Biology. 23, 487–493 (2016).

2. R. Ernst, R. Klemm, L. Schmitt, K. Kuchler, in Methods in Enzymology (Academic Press, 2005; http://www.sciencedi-rect.com/science/article/pii/S0076687905000261), vol. 400 of Phase II Conjugation Enzymes and Transport Systems, pp. 460–484.

3. E. Lamping, P. V. Baret, A. R. Holmes, B. C. Monk, A. Goffeau, R. D. Cannon, Fungal PDR transporters: Phylogeny, topology, motifs and function. Fungal Genetics and Biology. 47, 127–142 (2010).

4. G. Leppert, R. McDevitt, S. C. Falco, T. K. V. Dyk, M. B. Ficke, J. Golin, Cloning by gene amplification of two loci conferring multiple drug resistance in Saccharomyces. Genetics. 125, 13–20 (1990).

5. J. Golin, S. V. Ambudkar, The multidrug transporter Pdr5 on the 25th anniversary of its discovery: an important model for the study of asymmetric ABC transporters. Biochem J. 467, 353–363 (2015).

6. B. Rogers, A. Decottignies, M. Kolaczkowski, E. Carvajal, E. Balzi, A. Goffeau, The pleitropic drug ABC transporters from Saccharomyces cerevisiae. J. Mol. Microbiol. Biotechnol. 3, 207–214 (2001).

7. M. Kolaczkowski, van der R. Michel, A. Cybularz-Kolaczkowska, J.-P. Soumillion, W. N. Konings, G. André, Anticancer Drugs, Ionophoric Peptides, and Steroids as Substrates of the Yeast Multidrug Transporter Pdr5p. J. Biol. Chem. 271, 31543–31548 (1996).

8. M. Martínez, J. L. López-Ribot, W. R. Kirkpatrick, S. P. Bachmann, S. Perea, M. T. Ruesga, T. F. Patterson, Heterogeneous mechanisms of azole resistance in Candida albicans clinical isolates from an HIV-infected patient on continuous fluconazole therapy for oropharyngeal candidosis. J Antimicrob Chemother. 49, 515–524 (2002).

9. D. Sanglard, K. Kuchler, F. Ischer, J. L. Pagani, M. Monod, J. Bille, Mechanisms of resistance to azole antifungal agents in Candida albicans isolates from AIDS patients involve specific multidrug transporters. Antimicrobial Agents and Chemotherapy. 39, 2378–2386 (1995).

10. P. Leroux, R. Fritz, D. Debieu, C. Albertini, C. Lanen, J. Bach, M. Gredt, F. Chapeland, Mechanisms of resistance to fungicides in field strains of Botrytis cinerea. Pest Management Science. 58, 876–888 (2002).

11. L. Schmitt, R. Tampé, Structure and mechanism of ABC transporters. Current Opinion in Structural Biology. 12, 754–760 (2002).

12. T. Stockner, R. Gradisch, L. Schmitt, The role of the degenerate nucleotide binding site in type I ABC exporters. FEBS Letters. 594, 3815–3838 (2020).

13. R. P. Gupta, P. Kueppers, N. Hanekop, L. Schmitt, Generating Symmetry in the Asymmetric ATP-binding Cassette (ABC) Transporter Pdr5 from Saccharomyces cerevisiae. J. Biol. Chem. 289, 15272–15279 (2014).

14. M. Wagner, S. H. J. Smits, L. Schmitt, In vitro NTPase activity of highly purified Pdr5, a major yeast ABC multidrug transporter. Scientific Reports. 9, 7761 (2019).

15. Y. Cheng, Membrane protein structural biology in the era of single particle cryo-EM. Current Opinion in Structural Biology. 52, 58–63 (2018).

16. T. H. Bayburt, S. G. Sligar, Membrane Protein Assembly into Nanodiscs. FEBS Lett. 584, 1721–1727 (2010).

17. M. L. Carlson, J. W. Young, Z. Zhao, L. Fabre, D. Jun, J. Li, J. Li, H. S. Dhupar, I. Wason, A. T. Mills, J. T. Beatty, J. S. Klassen, I. Rouiller, F. Duong, The Peptidisc, a simple method for stabilizing membrane proteins in detergent-free solution. eLife. 7, e34085 (2018).

18. A. J. Rothnie, Detergent-Free Membrane Protein Purification. Methods Mol. Biol. 1432, 261–267 (2016).

19. J. Frauenfeld, R. Löving, J.-P. Armache, A. F.-P. Sonnen, F. Guettou, P. Moberg, L. Zhu, C. Jegerschöld, A. Flayhan, J. A. G. Briggs, H. Garoff, C. Löw, Y. Cheng, P. Nordlund, A saposin-lipoprotein nanoparticle system for membrane proteins. Nature Methods. 13, 345–351 (2016).

20. K. Rantalainen, Z. T. Berndsen, A. Antanasijevic, T. Schiffner, X. Zhang, W.-H. Lee, J. L. Torres, L. Zhang, A. Irimia, J. Copps, K. H. Zhou, Y. D. Kwon, W. H. Law, C. A. Schramm, R. Verardi, S. J. Krebs, P. D. Kwong, N. A. Doria-Rose, I. A. Wilson, M. B. Zwick, J. R. Yates, W. R. Schief, A. B. Ward, HIV-1 Envelope and MPER Antibody Structures in Lipid Assemblies. Cell Rep. 31, 107583 (2020).

21. Y.-C. Kuo, H. Chen, G. Shang, E. Uchikawa, H. Tian, X.-C. Bai, X. Zhang, Cryo-EM structure of the PlexinC1/A39R complex reveals inter-domain interactions critical for ligand-induced activation. Nature Communications. 11, 1953 (2020).

22. R. Ernst, P. Kueppers, C. M. Klein, T. Schwarzmueller, K. Kuchler, L. Schmitt, A mutation of the H-loop selectively affects rhodamine transport by the yeast multidrug ABC transporter Pdr5. PNAS. 105, 5069–5074 (2008).

23. R. Egner, F. E. Rosenthal, A. Kralli, D. Sanglard, K. Kuchler, Genetic Separation of FK506 Susceptibility and Drug Transport in the Yeast Pdr5 ATP-binding Cassette Multidrug Resistance Transporter. MBoC. 9, 523–543 (1998).

24. C. C. Goodno, in Methods in Enzymology (Academic Press, 1982; http://www.sciencedirect.com/science/arti-cle/pii/0076687982850143), vol. 85 of Structural and Contractile Proteins Part B: The Contractile Apparatus and the Cytoskeleton, pp. 116–123.

25. T. W. Loo, D. M. Clarke, Vanadate trapping of nucleotide at the ATP-binding sites of human multidrug resistance P-glycoprotein exposes different residues to the drug-binding site. PNAS. 99, 3511–3516 (2002).

26. J. Chen, S. Sharma, F. A. Quiocho, A. L. Davidson, Trapping the transition state of an ATP-binding cassette transporter: Evidence for a concerted mechanism of maltose transport. PNAS. 98, 1525–1530 (2001).

27. S. Hofmann, D. Januliene, A. R. Mehdipour, C. Thomas, E. Stefan, S. Brüchert, B. T. Kuhn, E. R. Geertsma, G. Hummer, R. Tampé, A. Moeller, Conformation space of a heterodimeric ABC exporter under turnover conditions. Nature. 571, 580–583 (2019).

28. S. H. W. Scheres, RELION: Implementation of a Bayesian approach to cryo-EM structure determination. Journal of Structural Biology. 180, 519–530 (2012).

29. P. B. Rosenthal, R. Henderson, Optimal Determination of Particle Orientation, Absolute Hand, and Contrast Loss in Single-particle Electron Cryomicroscopy. Journal of Molecular Biology. 333, 721–745 (2003).

30. J. Ye, A. R. Osborne, M. Groll, T. A. Rapoport, RecA-like motor ATPases—lessons from structures. Biochimica et Biophysica Acta (BBA)-Bioenergetics. 1659, 1–18 (2004).

31. R. M. Story, I. T. Weber, T. A. Steitz, The structure of the E. coli recA protein monomer and polymer. Nature. 355, 318–325 (1992).

32. C. Furman, J. Mehla, N. Ananthaswamy, N. Arya, B. Kulesh, I. Kovach, S. V. Ambudkar, J. Golin, The Deviant ATP-binding Site of the Multidrug Efflux Pump Pdr5 Plays an Active Role in the Transport Cycle. J. Biol. Chem. 288, 30420–30431 (2013).

33. N. M. I. Taylor, I. Manolaridis, S. M. Jackson, J. Kowal, H. Stahlberg, K. P. Locher, Structure of the human multidrug transporter ABCG2. Nature. 546, 504–509 (2017).

34. K. P. Locher, Structure and mechanism of ATP-binding cassette transporters. Philos Trans R Soc Lond B Biol Sci. 364, 239–245 (2009).

35. I. Manolaridis, S. M. Jackson, N. M. I. Taylor, J. Kowal, H. Stahlberg, K. P. Locher, Cryo-EM structures of a human ABCG2 mutant trapped in ATP-bound and substrate-bound states. Nature. 563, 426–430 (2018).

36. J. Peng, D. Schwartz, J. E. Elias, C. C. Thoreen, D. Cheng, G. Marsischky, J. Roelofs, D. Finley, S. P. Gygi, A proteomics approach to understanding protein ubiquitination. Nat. Biotechnol. 21, 921–926 (2003).

37. A. L. Hitchcock, K. Auld, S. P. Gygi, P. A. Silver, A subset of membrane-associated proteins is ubiquitinated in response to mutations in the endoplasmic reticulum degradation machinery. Proc. Natl. Acad. Sci. U.S.A. 100, 12735–12740 (2003).

38. J. M. Ren, T. Rejtar, L. Li, B. L. Karger, N-Glycan Structure Annotation of Glycopeptides Using a Linearized Glycan Structure Database (GlyDB). J Proteome Res. 6, 3162–3173 (2007).

39. I. L. Urbatsch, M. Julien, I. Carrier, M.-E. Rousseau, R. Cayrol, P. Gros, Mutational Analysis of Conserved Carboxylate Residues in the Nucleotide Binding Sites of P-Glycoprotein. Biochemistry. 39, 14138–14149 (2000).

40. C. Orelle, O. Dalmas, P. Gros, A. D. Pietro, J.-M. Jault, The Conserved Glutamate Residue Adjacent to the Walker-B Motif Is the Catalytic Base for ATP Hydrolysis in the ATP-binding Cassette Transporter BmrA. J. Biol. Chem. 278, 47002–47008 (2003).

41. P. Vergani, S. W. Lockless, A. C. Nairn, D. C. Gadsby, CFTR channel opening by ATP-driven tight dimerization of its nucleotide-binding domains. Nature. 433, 876–880 (2005).

42. D. C. Rees, E. Johnson, O. Lewinson, ABC transporters: The power to change. Nat Rev Mol Cell Biol. 10, 218–227 (2009).

43. J. Golin, Z. N. Kon, C.-P. Wu, J. Martello, L. Hanson, S. Supernavage, S. V. Ambudkar, Z. E. Sauna, Complete Inhibition of the Pdr5p Multidrug Efflux Pump ATPase Activity by Its Transport Substrate Clotrimazole Suggests that GTP as Well as ATP May Be Used as an Energy Source. Biochemistry. 46, 13109–13119 (2007).

44. J. Golin, S. V. Ambudkar, M. M. Gottesman, A. D. Habib, J. Sczepanski, W. Ziccardi, L. May, Studies with Novel Pdr5p Substrates Demonstrate a Strong Size Dependence for Xenobiotic Efflux. J. Biol. Chem. 278, 5963–5969 (2003).

45. S. M. Jackson, I. Manolaridis, J. Kowal, M. Zechner, N. M. I. Taylor, M. Bause, S. Bauer, R. Bartholomaeus, G. Bernhardt, B. Koenig, A. Buschauer, H. Stahlberg, K.-H. Altmann, K. P. Locher, Structural basis of small-molecule inhibition of human multidrug transporter ABCG2. Nat Struct Mol Biol. 25, 333–340 (2018).

46. B. J. Orlando, M. Liao, ABCG2 transports anticancer drugs via a closed-to-open switch. Nature Communications. 11, 2264 (2020).

47. D. Szöllősi, D. Rose-Sperling, U. A. Hellmich, T. Stockner, Comparison of mechanistic transport cycle models of ABC exporters. Biochimica et Biophysica Acta (BBA)-Biomembranes. 1860, 818–832 (2018).

48. M. L. Oldham, J. Chen, Snapshots of the maltose transporter during ATP hydrolysis. Proc. Natl. Acad. Sci. U.S.A. 108, 15152–15156 (2011).

49. M. Yang, N. Livnat Levanon, B. Acar, B. Aykac Fas, G. Masrati, J. Rose, N. Ben-Tal, T. Haliloglu, Y. Zhao, O. Lewinson, Single-molecule probing of the conformational homogeneity of the ABC transporter BtuCD. Nature Chemical Biology. 14, 715–722 (2018).

50. B. Verhalen, R. Dastvan, S. Thangapandian, Y. Peskova, H. A. Koteiche, R. K. Nakamoto, E. Tajkhorshid, H. S. Mchaourab, Energy transduction and alternating access of the mammalian ABC transporter P-glycoprotein. Nature. 543, 738–741 (2017).

51. N. Khunweeraphong, D. Szöllősi, T. Stockner, K. Kuchler, The ABCG2 multidrug transporter is a pump gated by a valve and an extracellular lid. Nature Communications. 10, 5433 (2019).

52. H. Osorio, E. Carvalho, M. del Valle, M. A. G. Sillero, P. Moradas-Ferreira, A. Sillero, H2O2, but not menadione, provokes a decrease in the ATP and an increase in the inosine levels in Saccharomyces cerevisiae. European Journal of Biochemistry. 270, 1578–1589 (2003).

53. A. Punjani, J. L. Rubinstein, D. J. Fleet, M. A. Brubaker, cryoSPARC: algorithms for rapid unsupervised cryo-EM structure determination. Nature Methods. 14, 290–296 (2017).

54. K. Zhang, Gctf: Real-time CTF determination and correction. Journal of Structural Biology. 193, 1–12 (2016).

55. J. Zivanov, T. Nakane, B. O. Forsberg, D. Kimanius, W. J. Hagen, E. Lindahl, S. H. Scheres, New tools for automated high-resolution cryo-EM structure determination in RELION-3. eLife. 7, e42166 (2018).

56. X. Bai, C. Yan, G. Yang, P. Lu, D. Ma, L. Sun, R. Zhou, S. H. W. Scheres, Y. Shi, An atomic structure of human γ-secretase. Nature. 525, 212–217 (2015).

57. S. H. W. Scheres, in Methods in Enzymology, R. A. Crowther, Ed. (Academic Press, 2016; http://www.sciencedi-rect.com/science/article/pii/S0076687916300301), vol. 579 of The Resolution Revolution: Recent Advances In cryoEM, pp. 125–157.

58. E. F. Pettersen, T. D. Goddard, C. C. Huang, G. S. Couch, D. M. Greenblatt, E. C. Meng, T. E. Ferrin, UCSF Chimera—A visualization system for exploratory research and analysis. Journal of Computational Chemistry. 25, 1605–1612 (2004).

59. A. Punjani, H. Zhang, D. J. Fleet, bioRxiv, in press, doi:10.1101/2019.12.15.877092.

60. J. Zivanov, T. Nakane, S. H. W. Scheres, Estimation of high-order aberrations and anisotropic magnification from cryo EM data sets in RELION-3.1. IUCrJ. 7, 253–267 (2020).

61. A. Rohou, N. Grigorieff, CTFFIND4: Fast and accurate defocus estimation from electron micrographs. Journal of Structural Biology. 192, 216–221 (2015).

62. K. Ramlaul, C. M. Palmer, C. H. S. Aylett, bioRxiv, in press, doi:10.1101/2019.12.12.874081.

63. K. Ramlaul, C. M. Palmer, C. H. S. Aylett, A Local Agreement Filtering Algorithm for Transmission EM Reconstructions. Journal of Structural Biology. 205, 30–40 (2019).

64. A. Leaver-Fay, M. Tyka, S. M. Lewis, O. F. Lange, J. Thompson, R. Jacak, K. Kaufman, P. D. Renfrew, C. A. Smith, W. Sheffler, I. W. Davis, S. Cooper, A. Treuille, D. J. Mandell, F. Richter, Y.-E. A. Ban, S. J. Fleishman, J. E. Corn, D. E. Kim, S. Lyskov, M. Berrondo, S. Mentzer, Z. Popović, J. J. Havranek, J. Karanicolas, R. Das, J. Meiler, T. Kortemme, J. J. Gray, B. Kuhlman, D. Baker, P. Bradley, ROSETTA3: an object-oriented software suite for the simulation and design of macromolecules. Methods Enzymol. 487, 545–574 (2011).

65. P. Emsley, K. Cowtan, Coot: model-building tools for molecular graphics. Acta Cryst D. 60, 2126–2132 (2004).

66. T. Burnley, C. M. Palmer, M. Winn, Recent developments in the CCP-EM software suite. Acta Cryst D. 73, 469–477 (2017).

67. T. I. Croll, ISOLDE: a physically realistic environment for model building into low-resolution electron-density maps. Acta Cryst D. 74, 519–530 (2018).

68. P. V. Afonine, B. K. Poon, R. J. Read, O. V. Sobolev, T. C. Terwilliger, A. Urzhumtsev, P. D. Adams, Real-space refinement in PHENIX for cryo-EM and crystallography. Acta Cryst D. 74, 531–544 (2018).

69. T. C. Terwilliger, S. J. Ludtke, R. J. Read, P. D. Adams, P. V. Afonine, Improvement of cryo-EM maps by density modification. bioRxiv, 845032 (2020).

70. A. J. Jakobi, M. Wilmanns, C. Sachse, Model-based local density sharpening of cryo-EM maps. eLife. 6, e27131 (2017).

71. T. D. Goddard, C. C. Huang, E. C. Meng, E. F. Pettersen, G. S. Couch, J. H. Morris, T. E. Ferrin, UCSF ChimeraX: Meeting modern challenges in visualization and analysis. Protein Sci. 27, 14–25 (2018).

72. R. A. Laskowski, M. B. Swindells, LigPlot+: Multiple Ligand–Protein Interaction Diagrams for Drug Discovery. J. Chem. Inf. Model. 51, 2778–2786 (2011).

73. E. Chovancova, A. Pavelka, P. Benes, O. Strnad, J. Brezovsky, B. Kozlikova, A. Gora, V. Sustr, M. Klvana, P. Medek, L. Biedermannova, J. Sochor, J. Damborsky, CAVER 3.0: a tool for the analysis of transport pathways in dynamic protein structures. PLoS Comput Biol. 8, e1002708 (2012).

74. Z. E. Sauna, S. S. Bohn, R. Rutledge, M. P. Dougherty, S. Cronin, L. May, D. Xia, S. V. Ambudkar, J. Golin, Mutations Define Cross-talk between the N-terminal Nucleotide-binding Domain and Transmembrane Helix-2 of the Yeast Multidrug Transporter Pdr5. J Biol Chem. 283, 35010–35022 (2008).

75. N. Ananthaswamy, R. Rutledge, Z. E. Sauna, S. V. Ambudkar, E. Dine, E. Nelson, D. Xia, J. Golin, The signaling interface of the yeast multidrug transporter Pdr5 adopts a cis conformation, and there are functional overlap and equivalence of the deviant and canonical Q-loop residues. Biochemistry. 49, 4440–4449 (2010).

76. P. Kueppers, R. P. Gupta, J. Stindt, S. H. J. Smits, L. Schmitt, Functional Impact of a Single Mutation within the Trans membrane Domain of the Multidrug ABC Transporter Pdr5. Biochemistry. 52, 2184–2195 (2013).

77. J. Mehla, R. Ernst, R. Moore, A. Wakschlag, M. K. Marquis, S. V. Ambudkar, J. Golin, Evidence for a molecular diode based mechanism in a multispecific ATP-binding cassette (ABC) exporter: SER-1368 as a gatekeeping residue in the yeast multidrug transporter Pdr5. J Biol Chem. 289, 26597–26606 (2014).

78. M. T. Downes, J. Mehla, N. Ananthaswamy, A. Wakschlag, M. Lamonde, E. Dine, S. V. Ambudkar, J. Golin, The Transmission Interface of the Saccharomyces cerevisiae Multidrug Transporter Pdr5: Val-656 Located in Intracellular Loop 2 Plays a Major Role in Drug Resistance. Antimicrob Agents Chemother. 57, 1025–1034 (2013).

